# A method to synthesize analytical rhodoquinone standards for quantitative analysis in tissue specimen

**DOI:** 10.64898/2026.05.04.722805

**Authors:** Thang Do, Akbar Ali, Jessica B. Spinelli

## Abstract

Rhodoquinone (RQ) is a recently discovered component of the mammalian electron transport chain (ETC) with a high degree of tissue-specificity. Currently, a lack of pure analytical standards limits efforts to precisely quantify its levels using liquid chromatography–tandem mass spectrometry (LC–MS/MS) and interrogate its biochemical functions within mammalian ETC complexes. Here, rhodoquinone-9 (RQ-9) and rhodoquinone-10 (RQ-10), and their isomeric by-products isorhodoquinone-9 (isoRQ-9) and isorhodoquinone-10 (isoRQ-10), were synthesized from ubiquinone-9 and ubiquinone-10 starting materials. Isomers were separated and purified by flash chromatography and structurally confirmed with nuclear magnetic resonance (NMR) spectroscopy. The chromatographic and fragmentation patterns of both the oxidized and reduced forms of these electron carriers were further characterized by LC–MS/MS, establishing signatures for their confident identification in lipidomics studies. LC-MS/MS analysis of murine kidney tissue with RQ-9 analytical standard spike-in corroborate the identity of the endogenous murine RQ-9 and enable absolute quantification of its levels. Thus, we synthesized and purified RQ-9 and RQ-10 analytical standards that will enable absolute quantification in mammalian tissues and *in vitro* reconstitution studies on RQ-9 and RQ-10 in the mammalian ETC.

## Introduction

The flow of electrons though electron transport chain (ETC) enables mitochondrial functions in bioenergetics, biosynthesis, balance of reduction and oxidation (redox) cofactors, cellular signaling, and epigenetic remodeling.^1, 2^ Ubiquinone (UQ), menaquinone (MQ), and rhodoquinone (RQ) are electron carriers that facilitate movement of electrons between the membrane-embedded ETC complexes, enabling these mitochondrial functions.^3^ The flow of electrons down this chain of reactions is dictated by the difference in the reduction potential (E°) of the electron donor and acceptor, with acceptors having higher E° values.^4, 5^ Ubiquinone (also known as Coenzyme Q) is “ubiquitous”, present in almost all organisms, and delivers its electrons to cytochrome C and ultimately oxygen (O_2_) as the terminal electron acceptor.^6, 7^ In contrast, MQ (also known as Vitamin K2) and RQ are species-specific and have lower E°, allowing them to deliver electrons to a broader range of terminal acceptors, including fumarate and nitrate.^3, 8^

Although RQ has long been thought to be restricted to bacteria,^9–15^ protists,^16^ parasitic helminths,^17, 18^ nematodes,^19, 20^ oysters, mussels, and snails,^3^ this metabolite has recently been found to be present in mammals, where it delivers electrons to fumarate, instead of O_2_, as the electron acceptor.^4^ In mammals, endogenous RQ is tissue-specific and much lower than UQ, indicating it is regulated in a cell-type and/or regional manner. In some bacteria, *C. elegans*, and in mice, the predominant form of RQ has a polyisoprene tail containing nine isoprene units (RQ-9);^7, 21^ whereas in humans and other bacterial strains, the major form has ten isoprene units (RQ-10).^22, 23^ As RQ can enable certain mitochondrial functions without the need for O_2_, the RQ/fumarate ETC has been explored for its therapeutic potential in protecting cells and tissues upon hypoxia exposure.^4^ Given the fundamental and translational implications of RQ, it is of utmost importance to be able to quantitatively measure this molecule in cells and tissues and perform *in vitro* reconsistution assays with it to elucidate its biochemical activities within the mammalian ETC.

Currently, there are no commercially available analytical standards for rhodoquinone, limiting the ability to perform quantitative analysis of its levels in biological materials by techniques such as targeted liquid chromatography−tandem mass spectrometry (LC−MS/MS). Although UV–Vis spectroscopy can be used to quantify RQ, it is less sensitive than LC–MS/MS and confounded by metabolites with overlapping absorption spectras. Beyond its use for quantitative analyses of endogenous RQ in cells and tissues, chemically pure RQ is necessary for reconstitution studies for structural determination of its binding pockets and kinetic parameters on ETC complexes. Previous attempts to synthesize, purify, and structurally validate RQ have been successful for a form that has a polyisoprene tail with three isoprene units (RQ-3).^24^ Although this compound is extremely useful for certain quantitative analyses, its shorter polyisoprene tail limits its ability to be used for docking and biochemical assays that recapitulate endogenous mammalian RQ. RQ-10 and iso-RQ10 have previously been synthesized from UQ-10; however, their structures were not confirmed definitively.^25, 26^

In stark contrast to the lack of chemically pure and structurally validated RQ-9 and RQ-10, UQ-9 and UQ-10 are commercially available, highly pure, inexpensive, and even widely sold as dietary supplements. Structurally, UQ and RQ are almost the same, differing primarily in one substituent on the quinone head group.^22^ This close structural similarity, particularly in the polyisoprene tail, makes UQ a practical precursor for the chemical synthesis and purification of RQ analytical standards. In this study, we sought to synthesize and purify rhodoquinone-9 and 10 analytical standards, enabling quantitative analysis of rhodoquinone in tissue specimen using LC-MS/MS.

## Materials and Methods

### Animals

7-week-old wild-type C57BL/6J mice (RRID:IMSR_JAX:000664) were bred and housed in the UMass Chan Medical School DiMare Center Animal Facility and maintained according to the Spinelli Lab protocol (PROTO202200067) approved by UMass Chan Medical School Institutional Animal Care and Use Committee (IACUC). At the time of tissue collection, mice were euthanized and immediately dissected to harvest both kidneys into an Eppendorf centrifuge tube. Collected tissues were snap-frozen in liquid nitrogen and stored in a -80°C freezer before further processing.

### RQ and isoRQ Synthesis from UQ Standards

Rhodoquinone (RQ) and isorhodoquinone (isoRQ) were synthesized from ubiquinone (UQ) by reaction with ammonia followed by purification by flash column chromatography. The reaction of UQ-9 or UQ-10 with aqueous ammonia was carried out in a mixture of diethyl ether and ethanol according to the reported method.^25, 26^ UQ-10 and UQ-9 were obtained from Ambeed. Ammonium hydroxide solution (28–30% ammonia) and ammonia solution in methanol (7 N) were purchased from Sigma-Aldrich and Fisher Scientific, respectively. All solvents (HPLC grade) were purchased from Sigma-Aldrich and used as received. Reactions were performed in oven-dried round bottomed flasks fitted with rubber septa. Automated flash column chromatography was performed on an Interchim PuriFlash system equipped with a UV–vis detector using prepacked silica gel cartridges from SiliCycle.

#### Methanolic ammonia Procedure

In a 150 mL round-bottom flask, UQ-9 or -10 (2.0 g) was dissolved in anhydrous diethyl ether (40 mL), giving a clear yellow-orange solution. The flask was flushed with argon and sealed with a rubber septum and parafilm. A solution of ammonia in methanol (∼7 N, 40 mL) was added, and the resulting reaction mixture was allowed to stand at room temperature for 72 hours. During this time, the color of the solution changed from orange to dark purple. The solvents were evaporated under reduced pressure on a rotary evaporator, keeping the bath temperature at 30 °C, to provide a dark brown viscid solid.

#### Aqueous ammonia Procedure

In a 150 mL round-bottom flask, UQ-9 or -10 (2.0 g) was dissolved in diethyl ether (25 mL), and ethanol (25 mL) was added to the clear orange solution. The flask was sealed with a rubber septum and parafilm. Ammonium hydroxide solution (28–30% ammonia solution in water, 2 mL) was added, and the reaction mixture was allowed to stand at room temperature for 5 days. During this time, the color of the solution changed from orange to dark purple. Solvents were evaporated under reduced pressure on a rotary evaporator, keeping the bath temperature at 30 °C. The residue was further dried under high vacuum to provide a dark brown viscid solid.

### RQ and isoRQ Purification

To purify the desired RQ and isoRQ products from this crude mixture, a three-step flash column chromatography method was developed. First, the two products were separated from the unreacted UQ and other by-products. For this, the crude reaction mixture was dissolved in a minimum volume of dichloromethane/hexanes (1:4) mixture and subjected to flash column chromatography using a silica gel column (SiliaSep, 80 g, equilibrated with hexanes, gradient elution with 0–25% diethyl ether/hexanes over 25 min). Fractions from this first purification run were analyzed by analytical TLC, which was performed using silica gel (60 F_254_) coated aluminum plates (EMD Millipore) with 75:25 hexanes: diethyl ether as eluent. In the reaction with UQ-10, it was found that about 1.0 g of starting material did not react. The fractions containing the purple quinone products were pooled and concentrated to provide a purple solid (0.25 g, ∼25% yield); TLC (25% diethyl ether/hexanes) indicated this sample contained a mixture of two products. According to literature reports, the upper and lower spots on the TLC correspond to RQ-10 and isoRQ-10, respectively.^26^ Next, the purple solid was dissolved in a minimum volume of dichloromethane/hexanes (1:5) mixture and again subjected to flash column chromatography using a silica gel column (SiliaSep Premium, 40 g, equilibrated with hexanes, gradient elution with 0–25% diethyl ether/hexanes over 25 min). The fractions containing the upper RQ-10 and lower isoRQ-10 products were pooled separately. Pure isoRQ-10 was obtained as a deep purple solid (45 mg); however, the RQ-10 sample was contaminated with small amount of isoRQ-10. Finally, the RQ-10 was purified by flash column chromatography using another silica gel column (SiliaSep Premium, 40 g, equilibrated with chloroform, elution with chloroform) to provide the pure RQ-10 as a purple solid (18 mg). Dried pure RQ and isoRQ standards were resuspended in 80:20 HPLC-grade EtOH (Sigma-Aldrich): hexanes (Fisher Scientific), then run and analyzed by the UQ/RQ C18 LC-MS Method.

### Pure RQ and isoRQ NMR Analysis

^1^H NMR (500 MHz, CDCl_3_), ^13^C NMR (126 MHz, CDCl_3_), DEPT-135 NMR (126 MHz, CDCl_3_), and 2D NMR spectra were acquired on a Bruker Avance III HD 500 MHz NMR instrument. The instrument probe temperature was set to 25 °C during each run. Chemical shifts are reported in ppm (δ scale) relative to the solvent signal and coupling constant (*J*) values are reported in hertz. Data are presented as follows: chemical shift, multiplicity (s = singlet, d = doublet, m = multiplet, dt = doublet of triplets), coupling constant in Hz, and integration. Structural assignments were made using a combination of 1D and 2D NMR spectra, including gradient selected COSY, HSQC, and HMBC experiments.

### Pure RQ and isoRQ Reduction

Vacuum dried RQ and isoRQ standards were resuspended in 80:20 HPLC-grade EtOH (Sigma-Aldrich): hexanes (Fisher Scientific) at a concentration that is detectible by LC-MS. To synthesize the reduced forms, 2 μL of 0.5 M sodium borohydride (Sigma-Aldrich) in 50 mM sodium hydroxide (Fisher Scientific) in HPLC-grade water (Sigma-Aldrich) was added to 50 μL of each standard. The mixture was vortexed briefly and let standing for about 10 minutes on dry ice. Samples were then run and analyzed by the UQ/RQ C18 LC-MS Method.

### Pure RQ Concentrations Measurement and Standard Calibration Curves

Pure RQ-9 and RQ-10 standards were resuspended in 80:20 HPLC-grade EtOH (Sigma-Aldrich): hexanes (Fisher Scientific). The standards concentrations were measured on the NanoDrop One (Thermo Scientific) by recording the absorbance at 283 nm for RQ-9 and for RQ-10, then calculated using the Beer-Lambert equation. A serial dilution of each standard was made at ratio 1 to 2, 5, 10, 25, 100, 1000, 5000, 10000, 50000, 100000, 1000000, and 10000000. Samples were then run and analyzed by the UQ/RQ C18 LC-MS Method to generate RQ-9 and RQ-10 calibration curves.

### RQ-10 Methanolic Solution and Light Exposure Stability Test

Concentrated RQ-10 standards resuspended in 80:20 HPLC-grade EtOH (Sigma-Aldrich): hexanes (Fisher Scientific) were diluted 1:100 in 100% LCMS-grade methanol (Sigma-Aldrich) with 2 mM HPLC-grade ammonium formate (Honeywell) to final concentration of 100 μM. Samples were then incubated in the dark (no light), under ambient light or UV light for 1 hour at room temperature. Levels of UQ-10 and RQ-10 were measured by running and analyzing samples using the UQ/RQ C18 LC-MS Method. Stability of RQ-10 was compared to standard in only 80:20 EtOH:hexanes with no light exposure.

### Mouse Tissue RQ-9 Extraction and Measurement

The extraction and detection of endogenous RQ-9 from mouse kidney on LC-MS was adapted from a published protocol.^27^ This protocol is briefly described below.

#### Mitochondrial Isolation from Mouse Kidney Tissues

Tissues were flash frozen in liquid nitrogen and powderized using a mortar and pestle. Approximately 50 mg of powder was moved to a 2.0 mL Eppendorf tube and resuspended in 2 mL of ice-cold mitochondria isolation buffer containing 200 mM sucrose (Sigma-Aldrich), 1 mM EGTA (Sigma-Aldrich), and 10 mM Tris (Fisher Scientific), adjusted to pH 7.4 using HEPES 1 M (Sigma-Aldrich) in Milli-Q water. The resuspended mixture was dounced 30 times using a PTFE tissue grinder (VWR) in a homogenizer on wet ice. The homogenates were centrifuged in another 2.0 mL Eppendorf tube at 600 × g for 10 minutes at 4 °C. Supernatant was moved to a new tube and centrifuged again at 7,000 × g for 10 minutes at 4 °C and then discarded. 2 mL of isolation buffer was added to wash the pellets and another 10-minute centrifugation at 7,000 × g and 4 °C was performed. The supernatant was discarded and the wash step above was repeated with 1mL of isolation buffer. Washed mitochondrial pellets were re-suspended in 1 mL of isolation buffer. A 5% volume (50 μL) aliquot of each sample was moved to a 1.5 mL Eppendorf tube before the spinning step for protein quantification to normalize mass spectrometry data. The remaining materials were stored in -80 °C until they were ready for UQ/RQ extraction.

#### Protein Quantification of Mitochondria Pellets

Aliquots of mitochondria in isolation buffer for protein quantification were pelleted by centrifugation at 7,000 × g for 10 minutes at 4 °C and then resuspended in 100 µL of 1:10 dilution of 10X RIPA Lysis Buffer (0.5M Tris-HCl, 1.5M NaCl, 2.5% deoxycholic acid, 10% NP-40, 10mM EDTA, pH 7.4) (EMD Millipore) diluted in MilliQ water and supplemented with one cOmplete EDTA-free protease inhibitor tablet (Roche) per 50 mL of buffer. Samples were vortexed for 10 minutes at 4 °C then clarified by centrifugation at 21,300 × g and 4 °C for 10 minutes. A Pierce BCA Protein Assay Kit (Life Technologies) was used to quantify protein content in the collected supernatant. The measured concentrations were extrapolated to calculate total mitochondrial protein in samples used for UQ/RQ isolation.

#### UQ/RQ Extraction from Isolated Mitochondria

The remaining 95% of the purified mitochondria were vortexed in ice-cold 500 μL 100% LCMS-grade methanol (Sigma-Aldrich) with 0.1% (w/v) HCl from 3 M hydrogen chloride solution in methanol (Sigma-Aldrich) for 10 minutes at 4 °C. Subsequently, 500 μL of 100% LCMS-grade hexanes (Fisher Scientific) was added to each sample and vortexed for an additional 10 minutes at 4 °C, followed by 10-minute centrifugation at 21,300 x g for 4 °C. The non-polar hexanes layer on top was transferred to a new Eppendorf tube. Samples were then moved a Refrigerated CentriVap Benchtop Vacuum Concentrator connected to a CentriVap-105 Cold Trap (Labconco) allowing the removal of solvent. Dried samples were stored in -80 °C freezer until ready to be resuspended and run on the LC-MS. With the total protein calculated from method described in section above, samples were resuspended in 100% LCMS-grade methanol (Sigma-Aldrich) with 2 mM HPLC-grade ammonium formate (Honeywell) at volumes that normalize samples to concentration of 10 μg mitochondrial protein per μL. In this protocol, the acidification of methanol:hexanes extraction buffers with methanolic HCl and resuspension buffer with ammonium formate helps stabilizing the redox status of the quinones and quinols extracted from mitochondria. Resuspended samples were then run and analyzed by the UQ/RQ C18 LC-MS Method.

### UQ/RQ C18 LC-MS Method

UQ and RQ levels were measured by the Q Exactive Orbitrap Mass Spectrometer consisting of a HESI II probe and Ion Max source coupled with a Vanquish Flex UHPLC system. The instruments underwent weekly clean and calibration with Pierce ESI Ion Calibration Calmix (Thermo Scientific) in both negative and positive ion mode. LC-MS samples were kept in a 4 °C autosampler before being injected into a Supelco Ascentis Express C18 (2.7µm) HPLC column (EMD Millipore). The column oven was maintained at 25 °C. The mobile phase composed of buffer A containing 60:40 HPLC-grade acetonitrile (Sigma-Aldrich): HPLC-grade water (Sigma-Aldrich) and buffer B containing 90:9:1 HPLC-grade 2-propanol (Sigma-Aldrich): HPLC-grade acetonitrile (Sigma-Aldrich): HPLC-grade water (Sigma-Aldrich). Both buffers were supplemented with 10 mM HPLC-grade ammonium formate (Honeywell) and 0.1% LC-MS grade formic acid (Fisher Scientific). The flow rate was maintained at 0.5 mL/min throughout each run. A linear gradient from 15% to 30% Buffer B was applied over the first 2.4 minutes. Buffer B was then increased from 30% to 48% between 2.4 and 3.0 minutes, followed by a gradual increase from 48% to 82% from 3.0 to 13.2 minutes. The composition was further increased from 82% to 99% Buffer B between 13.2 and 13.8 minutes and held at 99% from 13.8 to 15.4 minutes. Finally, Buffer B was decreased from 99% to 15% between 15.4 and 20.0 minutes. The mass spectrometer run was set at full scan ranged from 700 to 1200 m/z, 70,000 resolution, 3×10^6^ AGC target, and 200 msec of maximum injection time. The HESI source was operated with a sheath gas flow of 40 units, an auxiliary gas flow of 15 units, and a sweep gas flow of 1 unit. The spray voltage was 4 kV, with the capillary temperature maintained at 320 °C and the auxiliary gas heater temperature set to 30 °C.

To obtain fragmentation patterns, a product reaction monitoring (PRM) was included in the method to detect positive ions (RQ-9 and isoRQ-9: m/z 780.6294, RQH_2_-9 and isoRQH_2_-9: m/z 782.6451, RQ-10 and isoRQ-10: m/z 848.6921, RQH_2_-10 and isoRQH_2_-10: m/z 850.7077, all masses are for proton adducts) at resolution of 17,500, AGC target of 2 x10^5^. The PRM maximum IT was set to 150 ms, isolation window to 0.5 m/z, and Stepped Normalized Collision Energy (Stepped NCE) to 30. (Iso)rhodoquinone-9, (iso)rhodoquinol-9, (iso)rhodoquinone-10, (iso)rhodoquinol-10 m/zs for proton adducts were targeted in the PRM inclusion list with monitoring window of 12-15-minute, 12-15-minute, 13-16-minute, and 13-16-minute respectively.

### Absolute Quantification of Endogenous RQ with Pure Synthesized Standard

Resuspended rhodoquinone-9 extracted from mouse kidney was made into two aliquots equal in volume. One aliquot was spiked in rhodoquinone-9 standard to reach a final concentration of 1 μM for synthetic RQ-9, the other was added with an equivalent volume of 80:20 HPLC-grade EtOH (Sigma-Aldrich): hexanes (Fisher Scientific) as solvent control. Both samples were then run and analyzed by the UQ/RQ C18 LC-MS Method. Concentration of endogenous rhodoquinone-9 in the resuspended lysate was calculated by dividing the RQ-9 ion counts from non-spiked sample by the difference between spiked and non-spiked then multiplying this ratio by the final concentration of the spiked analytical RQ-9 standard (1 μM).

### Chromatographic Resolution between RQ-9/isoRQ-9 and Endogenous Aminoquinones

To confirm that the endogenous aminoquinone detected in mouse samples is RQ-9; on the same LC-MS analytical run, pure RQ-9 and isoRQ-9 standards prepared in 80:20 (v/v) HPLC-grade ethanol (Sigma-Aldrich): hexanes (Fisher Scientific), along with a sample of rhodoquinone-9 extracted from mouse kidney mitochondria were run and analyzed using UQ/RQ C30 LC-MS Method. To further confirm peak identity by co-elution, RQ-9 and isoRQ-9 standards were spiked separately into two independent aliquots of the mouse kidney mitochondrial lysate. Spiked samples were briefly vortexed to ensure homogeneous mixing before LC-MS analysis using the same UQ/RQ C30 Method.

### UQ/RQ C30 LC-MS Method

Similar to the UQ/RQ C18 LC-MS Method, the mass spectrometry was performed on the Q Exactive Orbitrap Mass Spectrometer consisting of a HESI II probe and Ion Max source coupled with a Vanquish Flex UHPLC system. Samples were kept in a 4 °C autosampler before being injected into a Supelco Ascentis Express 160 Å C30 (2.7µm) HPLC column (EMD Millipore). The column chamber was maintained at 50 °C. The mobile phase composed of buffer A containing 60:40 HPLC-grade acetonitrile (Sigma-Aldrich): HPLC-grade water (Sigma-Aldrich) and buffer B containing 88:10:2 HPLC-grade 2-propanol (Sigma-Aldrich): HPLC-grade acetonitrile (Sigma-Aldrich): HPLC-grade water (Sigma-Aldrich). Both buffers were supplemented with 10 mM HPLC-grade ammonium formate (Honeywell) and 0.1% LC-MS grade formic acid (Fisher Scientific). The flow rate was maintained at 0.4 mL/min throughout with a three-minute equilibration at 30% Buffer B prior to each run. This initial percentage of Buffer B was maitained during the first minute, followed by an increase to 45% for the next two minutes. A linear gradient from 45% to 75% Buffer B was applied over the next 10.0 minutes. Buffer B was then increased from 75% to 90% between 13.0 and 14.5 minutes, followed by a gradual increase from 90% to 95% from 14.5 to 20.0 minutes. The composition was further increased from 95% to 100% Buffer B between 20.0 and 20.1 minutes and held at 100% from 20.1 to 25.0 minutes. Finally, Buffer B was decreased from 100% to 30% between 25.0 and 25.1 minutes, then remained at 30% until 28 minutes. The mass spectrometer run was set at full scan ranged from 700 to 1200 m/z, 70,000 resolution, 3×10^6^ AGC target, and 200 msec of maximum injection time. The HESI source was operated with a sheath gas flow of 40 units, an auxiliary gas flow of 15 units, and a sweep gas flow of 1 unit. The spray voltage was 4 kV, with the capillary temperature maintained at 320 °C and the auxiliary gas heater temperature set to 30 °C.

### Data Analysis and Visualization

Extracted ions chromatograms (EICs) and MS^2^ fragmentation spectra were viewed on Xcalibur Qual Browser (Thermo Scientific) and exported to Microsoft Excel for external plotting. Peak areas for endogenous rhodoquinone-9, rhodoquinone-9 and - 10 standards were analyzed on TraceFinder 5.1 (ThermoFisher Scientific). Peaks were selected based on their retention time and fragmentation patterns consistent with pure synthesized standards. Peaks integration for ions count measurement adhered strictly to the 5 ppm mass tolerance, integrated adducts are listed either on the main text or figure legends. Simple linear regression was performed in Microsoft Excel with the calibration curve constrained to pass through the origin. The lower limit of detection (LOD) and limit of quantification (LOQ) were calculated from the regression parameters (the slope and standard deviation of the curve) derived from the three lowest calibration points exhibiting a visible signal in the EICs, following the referenced protocol.^28^ The statistical tests were performed on GraphPad Prism version 10.6.1 and are stated in the figure legends where applicable. N values are included as number of technical repliactes and a p-value of 0.05 as threshold for statistical significant. All data was visualized using GraphPad Prism version 10.6.1, and relevant regression parameters were reported where applicable. Chemical structures were sketched and annotated in ChemDraw 23.1.2 (Revvity Signals). NMR data was processed and analyzed using TopSpin 4.5.0 (Bruker).

## Results

### Ammonolysis of ubiquinone generates a mixture of rhodoquinone and isorhodoquinone

As electron carriers, both UQ and RQ contain two para-oriented carbonyl groups at the C-1 and C-4 positions of the quinone ring, which enables reversible redox cycling. Both molecules share a hydrophobic polyisoprene tail extending from the C-5 position and a methyl substituent at the C-6 position, features that facilitate membrane localization and mobility within the mitochondrial lipid bilayer.^7, 21^ UQ possesses two methoxy groups on carbons C-2 and C-3 of the quinone ring while RQ has an amino group on carbon C-2 and a methoxy group on carbon C-3.^22^ Given the structural similarities of these molecules, RQ with 9 and 10 isoprene units can be synthesized from the corresponding UQ-9 and UQ-10 by reaction with aqueous ammonia, as previously reported.^25, 26^ However, the separation of RQ10 and isoRQ10 required repeated preparative thin-layer chromatography (TLC) using the multiple development procedure. ^26^

Methoxy groups in coenzyme Q are viable leaving groups due to high electrophilicity of the quinone ring, making UQ susceptible to nucleophilic substitution in the presence of ammonia.^25^ Consistent with previous findings, methanolic or aqueous ammonia were incubated with UQ-9 or UQ-10, producing an isomeric mixture of rhodoquinone-9 or -10 (RQ-9 or -10, 2-amino-3-methoxy-5-methyl-6-(nona/deca)prenyl-1,4-benzoquinone) where the 2-methoxy group was substituted by an amino group and isorhodoquinone-9 or -10 (isoRQ-9 or -10, 3-amino-2-methoxy-5-methyl-6-(nona/deca)prenyl-1,4-benzoquinone) where the 3-methoxy group was substituted by an amino group (Figure 1A).^25, 26^ This reaction changed the solution color from a bright yellow, indicative of UQ, to an earthy brown mixture as combination of the residual starting material and the deep purple color that is a characteristic of RQ.

**Figure 1.**
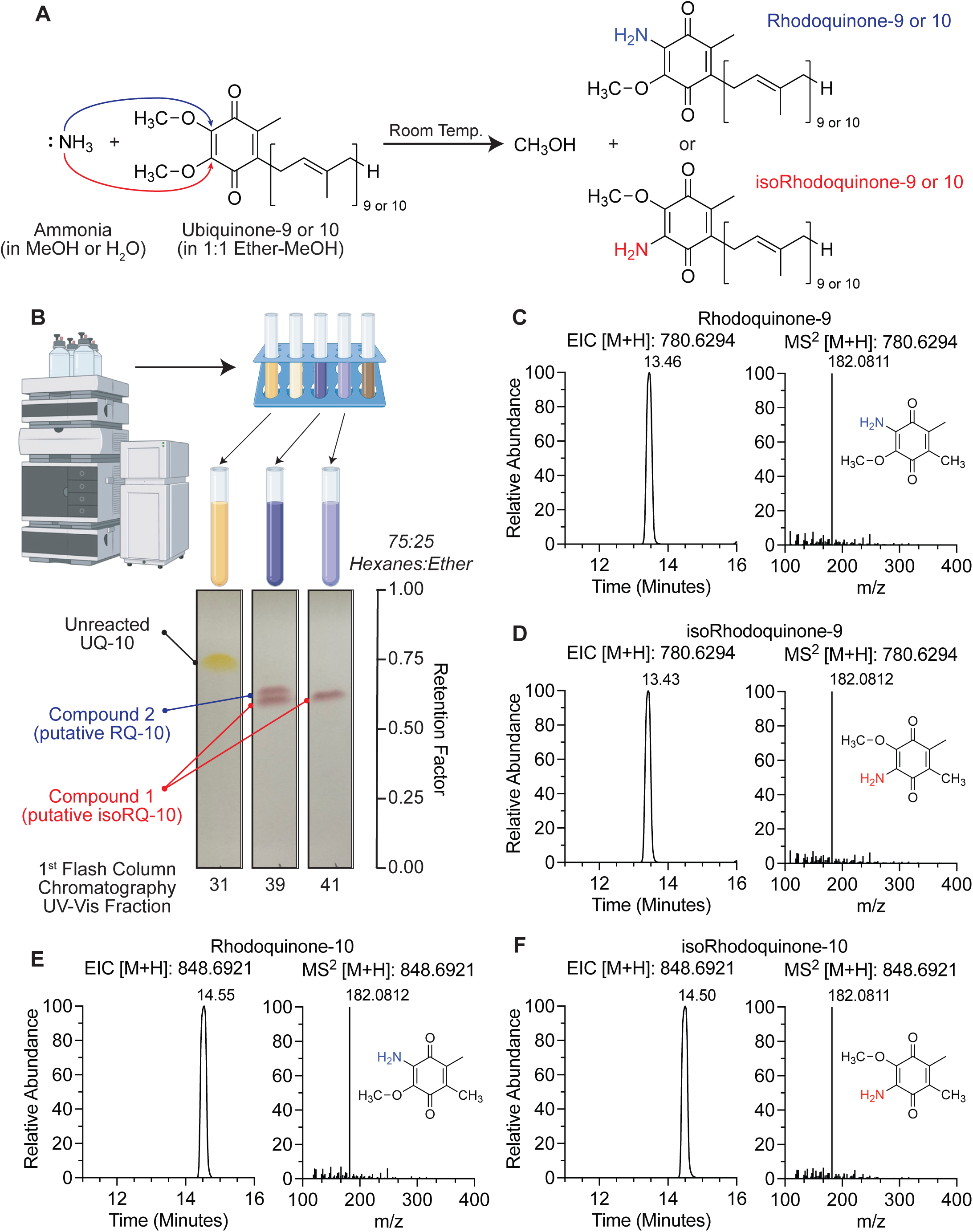
Synthesis of rhodoquinone and isorhodoquinone analytical standards from ubiquinone. (A) Schematic of reaction to synthesize rhodoquinone and isorhodoquinone from ubiquinone with aqueous or methanolic ammonia. (B) Thin-Layer Chromatography (TLC) analysis of flash column chromatography UV-Vis fractions of crude product from rhodoquinone-10 and isorhodoquinone-10 synthesis reaction after 5 days with 75:25 hexanes: diethyl ether as eluent. (C – F) Liquid Chromatography Tandem Mass Spectrometry (LC-MS/MS) analysis of purified rhodoquinone-9, isorhodoquinone-9, rhodoquinone-10, and isorhodoquinone-10 collected in positive ion mode. Parent and product m/zs reported as the proton adduct.

To separate putative RQ and isoRQ species in this reaction, repeated purifications using flash column chromatography were performed. Briefly, dry crude product was resuspended a minimum volume of 1:4 dichloromethane: hexanes. The resulting mixture was purified by flash column chromatography on silica gel (SiliaSep, 80 g) using a hexanes/diethyl ether gradient (0–25%) over 25 minutes on an Interchim PuriFlash system equipped with a UV–vis detector. Fractions collected after the first flash column chromatography run from UQ-10 reaction, which did not yet separate the isomers, were analyzed by thin-layer chromatography (TLC) using silica gel (60 F_254_) coated aluminum plates with 75:25 hexanes: diethyl ether as the solvent system. TLC analysis showed that some fractions contained a mixture of two purple products which, according to literature reports, reflect isoRQ as the lower spot (purple **Compound 1**) and RQ as the upper spot (purple **Compound 2**).^26^ Retention factor R_f_ values for **Compound 1** and **2** were 0.60 and 0.62 respectively, both are lower than UQ at 0.74 (Figure 1B). The purple fractions containing putative quinone products were pooled and concentrated to provide a purple solid (0.25 g). To separate these two putative isomers, two more rounds of flash chromatography were performed on the pooled purple fractions. Consistent with previous reports from the reaction with UQ-10, **Compound 1** was obtained as the major product (about 45 mg) and **Compound 2** as the minor product (about 18 mg) after purification.

We next sought to validate that the molecules we synthesized and purified are RQ isomers. To this end, the pure compounds were resuspended in 80:20 HPLC-grade ethanol: hexanes and analyzed with LC-MS/MS using a C18 column. Extracted ions chromatograms were taken for the proton adducts [M+H]^+^ collected in positive ion mode for RQ-9 and isoRQ-9 with *m/z* 780.6294, and RQ-10 and isoRQ-10 with m/z 848.6921. The mass tolerance thresholds utilized for generating these extracted ion chromatograms were ±5 ppm. We observed only a slight difference in retention time between the **Compound 1** (putative isoRQ) and **Compound 2** (putative RQ). Accordingly, the putative RQ species eluted slightly after the putative isoRQ (Figure 1C-F), and both eluted prior to UQ. This order of elution is consistent with TLC R_f_ values (Figure 1B). To further interrogate the identity of these putative RQ and isoRQ standards, we performed MS^2^ analysis and compared it to the well documented fragmentation signature of RQ-9 and RQ-10.^4^ All isomers containing either 9-unit or 10-unit polyisoprene chain had the same fragmentation patterns with the characteristic fragment corresponding to the quinone head portion of the molecule at *m/z* 182.0811–182.0812 (Figure 1C–F). This result increased our confidence that we successfully synthesized and purified RQ, however, these data do not resolve which of the two isomers that we generated in the reaction is the endogenous form of RQ present in mammals.

### Nuclear Magnetic Resonance (NMR) characterization of purified rhodoquinone-10 and isorhodoquinone-10 confirm position of the amino group in each isomer

Because putative RQ and isoRQ could not be distinguished based on fragmentation properties, ^1^H, ^13^C, DEPT-135, and 2D NMR spectra (COSY, HSQC, and HMBC) were acquired on both purified standards to unambiguously assign the position of the amino substituent on the quinone headgroup. Overall, the ^13^C NMR spectra of RQ and isoRQ are similar except for the signals corresponding to the head groups (Figure 2A–B, S2). As expected, analysis of the ^13^C NMR resonances that correspond to the carbons on the poly-isoprene tail overlap closely between the two isomers generated in the reaction. The most notable differences are observed in the quinone headgroup region, with the chemical shifts associated with C-1 and C-4 carbonyl carbons flipped between the two isomers. The more downfield resonance corresponds to the carbonyl carbon adjacent to the carbon bearing the amino group (C-4 for **Compound 1** and C-1 for **Compound 2**) (Figure 2A–B).

**Figure 2.**
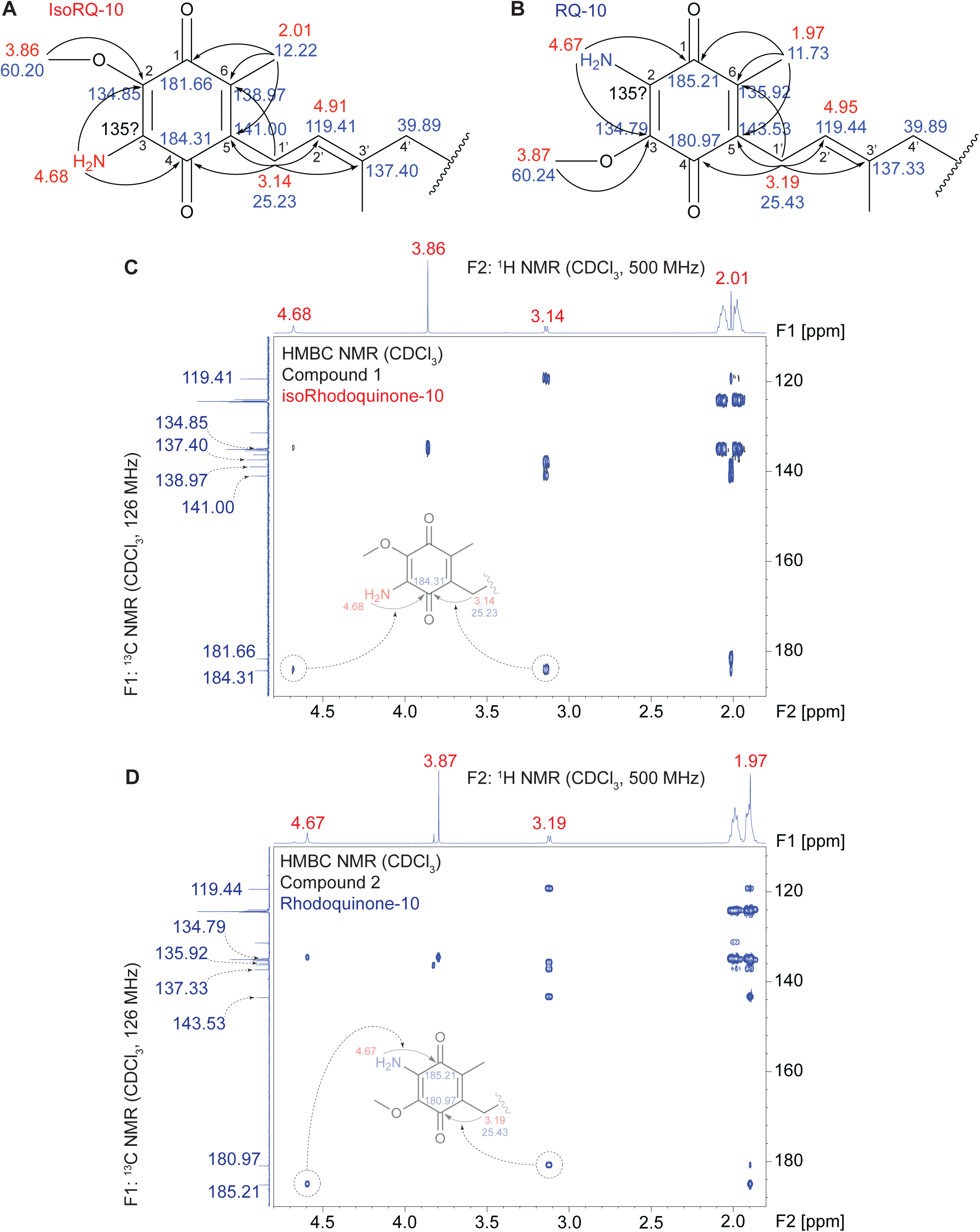
Nuclear Magnetic Resonance (NMR) analysis of purified rhodoquinone-10 and isorhodoquinone-10 confirms position of the amino group in each isomer. (A – B) Schematic of proton (^1^H) and carbon 13 (^13^C) chemical shift assignments with long-range correlation between carbons and hydrogens around the quinone head region of isorhodoquinone-10 and rhodoquinone-10. (C – D) Heteronuclear Multiple Bond Correlation (HMBC) NMR analysis of the quinone ring region in isorhodoquinone-10 and rhodoquinone-10. Highlighted correlations depict the positional difference of the amino group on isorhodoquinone and rhodoquinone. Also see Figures S1 – S10.

#### Compound 1

^1^H NMR (500 MHz, CDCl_3_): δ 5.13–5.05 (m, 9H), 4.94 (dt, *J* = 7.0, 1.0 Hz, 1H, H-2′), 4.68 (s, 2H, NH_2_), 3.86 (s, 3H, OCH_3_), 3.14 (d, *J* = 7.10 Hz, 2H, H-1′), 2.09–2.02 (m, 18H), 2.01 (s, 3H, CH_3_ at C-6), 1.99–1.94 (m, 18H), 1.73 (s, 3H, CH_3_ at C-3′), 1.68 (d, *J* = 1.0 Hz, 3H), 1.60 (s, 24H), 1.58 (s, 3H) ppm (Figure 2A–B, S1). ^13^C NMR (126 MHz, CDCl_3_): δ 184.31 (*C*=O, C-4), 181.66 (*C*=O, C-1), 141.00 (C-5), 138.97 (C-6), 137.40 (C-3′), 136.29 (C), 135.38 (C), 135.13 (C), 135.10 (C), 135.06 (C), 135.03 (C), 134.85 (C-3), 131.39 (C), 124.55 (CH), 124.41 (CH), 124.32 (CH), 124.01 (CH), 119.41 (CH-2′), 60.20 (O*C*H_3_ at C-2), 39.89 (CH_2_), 39.87 (CH_2_), 39.84 (CH_2_), 26.91 (CH_2_), 26.87 (CH_2_), 26.82 (CH_2_), 26.68 (CH_2_), 25.84 (CH_3_), 25.23 (CH_2_-1′), 17.83 (CH_3_), 16.46 (CH_3_ at C-3′), 16.17 (CH_3_), 12.22 (CH_3_ at C-6) ppm (Figure 2A–B, S2).

#### Compound 2

^1^H NMR (500 MHz, CDCl_3_): δ 5.13–5.05 (m, 9H), 4.95 (dt, *J* = 7.0, 1.1 Hz, 1H, H-2′), 4.67 (s, 2H, NH_2_), 3.87 (s, 3H, OCH_3_), 3.19 (d, *J* = 7.10 Hz, 1H, H-1′), 2.10–2.02 (m, 18H), 1.99–1.93 (m, 18H), 1.97 (s, 3H, overlapping, CH_3_ at C-6), 1.74 (s, 3H, CH_3_ at C-3′), 1.68 (d, *J* = 1.0 Hz, 3H), 1.60 (s, 24H), 1.58 (s, 3H) ppm (Figure 2A–B, S6). ^13^C NMR (126 MHz, CDCl_3_): δ **185.21** (*C*=O, C-1), **180.97** (*C*=O, C-4), **143.53** (C-5), **137.33** (C-3′), 136.22 (C), **135.92** (C-6), 135.34 (C), 135.12 (C), 135.08 (C), 135.05 (C), 135.03 (C), **134.79** (C-3), 131.39 (C), 124.55 (CH), 124.40 (CH), 124.32 (CH), 123.97 (CH), 119.44 (CH-2′), 60.24 (O*C*H_3_ at C-2), 39.89 (CH_2_), 39.87 (CH_2_), 26.91 (CH_2_), 26.86 (CH_2_), 26.82 (CH_2_), 26.70 (CH_2_), 25.84 (CH_3_), 25.43 (CH_2_-1′), 17.83 (CH_3_), 16.50 (CH_3_ at C-3′), 16.17 (CH_3_), 11.73 (CH_3_ at C-6) ppm (Figure 2A–B, S7).

The ^13^C NMR chemical shifts for the head group of **Compound 2** closely match with that of a previously synthesized analog of RQ-10 that has 3 isoprene units (RQ-3).^24^ Most notable chemical shifts in RQ-3 overlapping with the putative RQ-10 include δ **185.1** and **180.8** ppm for the two carbonyl carbons; δ **143.4**, **137.2**, **135.8**, and **134.7** ppm corresponding to C-5, C-3’, C-6, and C-3 respectively.^24^ Thus far, these analyses point to the assignments of **Compound 1** as isoRQ-10 and **Compound 2** as RQ-10.

Two other key assignments were the protons of the amino group and the benzylic methylene (H-1′) in the first isoprene unit directly attached to the aromatic ring at C-5. The NH_2_ and H-1′ chemical shifts in each compound were confirmed using a combination of 1D NMR (1H, 13C, DEPT-135) (Figure S1–S3, S6–S8) and 2D NMR (COSY, HSQC, HMBC) spectra (Figure S4–S5, S9–S10).

Finally, the assignment of the amino group position on the quinone ring was confirmed by HMBC correlations of key NMR signals. The observed HMBC correlations of the amino (NH_2_), methyl (at C-6), and benzylic methylene (H-1′) protons were analyzed, especially with the two carbonyl carbons at the C1 and C4 positions. In RQ-10, where the amino group occupies the C-2 position, the NH protons and the benzylic methylene protons do not correlate with the same carbonyl group. The NH_2_ protons correlate with the C-1 carbonyl, and the methylene protons correlate with the C-4 carbonyl, thus indicating that these two moieties are at opposing ends of the quinone head. In contrast, for isoRQ-10 which should have the amino group at the C-3 position, both the NH protons and the C-5 benzylic methylene protons exhibit long-range correlations to the same carbonyl carbon (C-4) (Figure 2A–D, S5, S10). Thus, we definitively identified which isomer is RQ and isoRQ and confirmed successful synthesis and purification of these two structural isomers.

### Chemical reduction of rhodoquinone and isorhodoquinone with sodium borohydride enables LC–MS characterization of corresponding quinols

In its physiological role as an electron carrier for the ETC, RQ toggles between its reduced and oxidized states. Similar to UQ, the redox state of RQ serves as an important indicator of ETC-linked mitochondrial functions. Therefore, the ability to identify and quantify rhodoquinol (RQH_2_), the reduced form of RQ, is critical. However, no analytical standards are currently available for rhodoquinol-9 or -10 to characterize their chromatographic and fragmentation properties on the LC-MS/MS. To generate and characterize rhodoquinol and isorhodoquinol on the mass spec, we shifted the redox equilibrium of RQ-9, RQ-10, isoRQ-9, and isoRQ-10 standards toward the reduced state by treatment with sodium borohydride (0.5 M in 50 mM NaOH) (Figure 3A). Samples were mixed by brief vortex then immediately analyzed on LC-MS with our C18 UQ/RQ method to avoid re-oxidation. Notably, tissue samples processed for RQ and RQH_2_ detection are extracted in acidified buffers using previously described methodology,^29^ which stabilizes the endogenous pool of reduced quinones. We observed similar stabilization of RQH_2_ when comparing standards resuspended in an acidic methanol ammonium formate solution versus a neutral ethanol:hexanes mixture (Figure S11 A-D). This stability of the reduced state was further unaffected by exposure to ambient or UV light (Figure S11A-D). However, due to the susceptibility of quinones to photodegradation upon UV absorption,^30^ it is the best practice to store resuspended standards long-term in the -80 °C with minimal light exposure to ensure accurate concentrations and preservation of redox states for LC-MS analysis.

**Figure 3.**
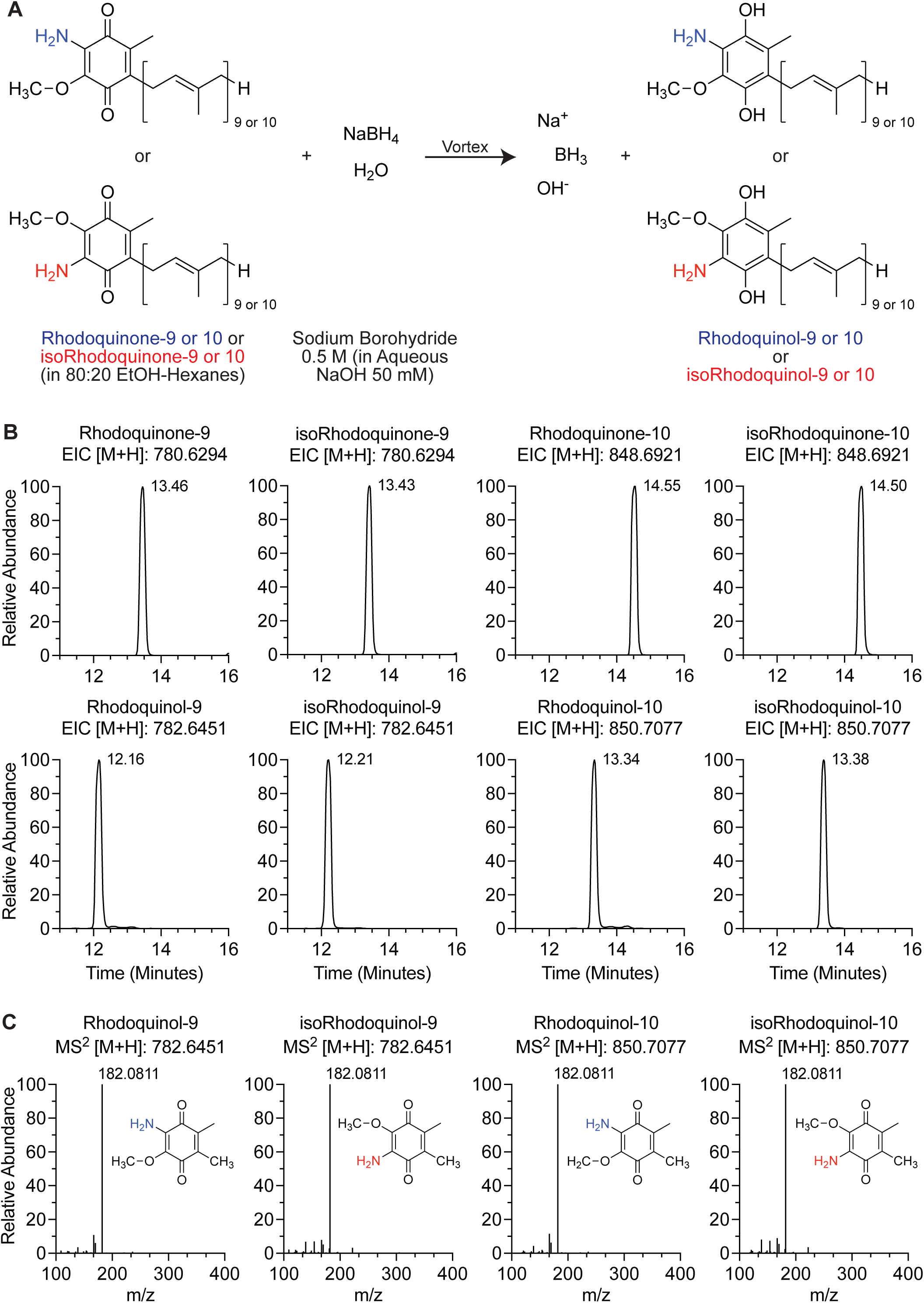
Reduction of rhodoquinone and isorhodoquinone by sodium borohydride reveals chromatographic properties and fragmentation patterns of rhodoquinol and isorhodoquinol. (A) Schematic of reaction to reduce pure rhodoquinone and isorhodoquinone analytical standards with sodium borohydride in basic solution. (B) Chromatograms of rhodoquinol-9, isorhodoquinol-9, rhodoquinol-10, and isorhodoquinol-10 compared to their respective oxidized precursors indicating upshifts in retention time of reduced forms on the chromatography. (C) Fragmentation patterns of rhodoquinol-9, isorhodoquinol-9, rhodoquinol-10, and isorhodoquinol-10 at their respective retention times.

For analysis of the quinols synthesized from the reaction, extracted ions chromatograms (EICs) and MS^2^ fragmentation spectra of RQH_2_-9 and isoRQH_2_-9 were obtained with positive proton adduct ions [M+H]^+^ at *m/z* of 782.6451 and at *m/z* of 850.7077 for RQH_2_-10 and isoRQH_2_-10 with a strict ±5 ppm tolerance. Consistent with the more polar nature of quinones upon reduction, each isomer of rhodoquinol eluted earlier on the chromatography than their respective oxidized forms (Figure 3B). This is consistent with the retention time differences of UQ and UQH_2_.^31^ RQH_2_-9 and 10 elute about 1.3 minutes before RQ-9 and 10 while isoRQH_2_-9 and 10 elute about 1.2 minutes before isoRQ-9 and 10 (Figure 3B), providing a subtle distinguishing feature between the two isomers. Moreover, the quinols exhibited the same fragmentation pattern as the quinones, with leading fragment of *m/z* 182.0811–182.0812 (Figure 3C). This fragmentation signature occurs because of oxidation of the quinol head group in the collision chamber during fragmentation, reverting it back to the quinone state. This phenomenon is also true of ubiquinol during LC-MS/MS fragmentation, which gave the same leading fragment as the oxidized state.^31^ Together, these chromatographic and fragmentation patterns provide reliable analytical features for identifying rhodoquinol in biological samples and enable their accurate detection and quantification by LC–MS/MS.

### Absolute quantification of endogenous rhodoquinone-9 in mouse kidney by LC–MS using the synthetic analytical RQ standard

In metabolomics studies, pure analytical standards serve to de-orphan newly identified metabolites by their mass to charge ratio *m/z*, ionization behaviors, retention times, and fragmentation spectra. Another critical use of chemically pure standards is absolute quantification of endogenous levels of the metabolite of interest. This is achieved by running a standard curve on the LC-MS setup and spiking-in pure analytical standard into the biological material to account for matrix effects on ionization efficiency of the endogenous metabolites.^32, 33^ As RQ is a newly identified component of mammalian mitochondria, we sought to use our purified RQ analytical standards to accurately quantify RQ levels in mouse kidney. First, the lower limit of detection (LOD) and limit of quantification (LOQ) was determined for RQ-9 and RQ-10 analytical standards. A calibration curve of each standard was obtained by running serial diluted samples up to very low concentrations and the LOD and LOQ were calculated using a reported method,^28^ involving parameters of linear regression analysis of lowest three data points where signals are clearly visible and mass accurate on the EICs. From resulting calibration curves, LOD and LOQ were estimated to be 82 nM and 248 nM, respectively, for RQ-9, and 46 nM and 141 nM, respectively, for RQ-10 (Figure 4A–B).

**Figure 4.**
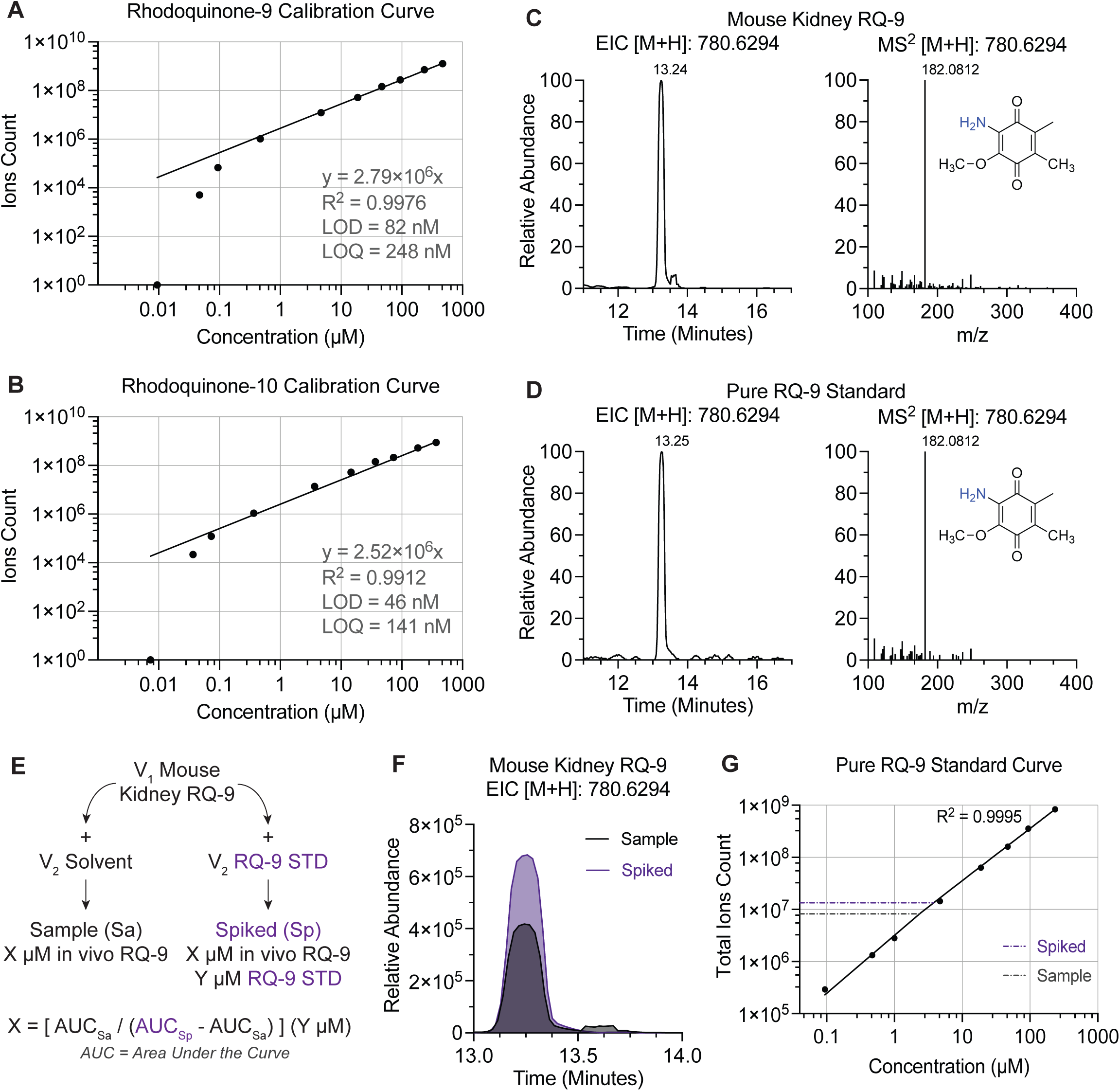
Absolute quantification of endogenous rhodoquinone-9 in mouse kidney tissue with analytical rhodoquinone-9 standard by LC-MS. (A – B) Rhodoquinone-9 and rhodoquinone-10 calibration curves from pure synthesized analytical standards with calculated lower limit of detection (LOD) and lower limit of quantification (LQD). (C – D) Chromatograms and fragmentation patterns of rhodoquinone-9 extracted from mouse kidney purified mitochondria compared to pure synthesized rhodoquinone-9 analytical standard. (E) Sample preparation and workflow to calculate the endogenous concentration of rhodoquinone-9 in mouse kidney tissue using LC-MS. (F) Overlay of proton adduct chromatograms for rhodoquinone-9 in mouse kidney sample alone and with pure standard spike-in. (G) Rhodoquinone-9 standard curve for linear range verification in absolute quantification of mouse kidney rhodoquinone-9. Total RQ-9 ions count consisting of the following adducts: proton [M+H], double proton [M+2H], sodium [M+Na], potassium [M+K], ammonium [M+NH_4_], and copper adducts [M+Cu]. Lines depicting the levels of RQ-9 in the kidney sample and the levels of RQ-9 in the kidney sample analytical standard spike in.

Next, the chromatographic properties of RQ-9 *in vivo* were compared to that of the pure analytical standard to confirm its identity and structure. Endogenous RQ-9 was extracted from mitochondria isolated from murine kidney tissue and LC-MS parameters were compared to the purified standard.^27^ Consistent with the endogenous kidney RQ-9 and synthetic RQ-9 standard being the same molecule, the EIC [M+H]^+^ showed the same retention time (RT) for both (Figure 4C–D). Moreover, the dominant fragment at *m/z* 182.0811–182.0812 was observed in the aminoquinone detected in kidney mitochondria and overlays exactly with the analytical RQ-9 standard we synthesized (Figure 4C–D). Since there was no clear separation between RQ-9 and isoRQ-9 on the C18 chromatography, we also analyzed these pure standards, kidney samples, and samples with spiked-in standards on a C30 chomatography method to provide better resolution of isomers and identify which isomer is dominant in kidney. The ovelaid EICs showed a slight backshift of pure isoRQ-9 peaks compared to pure RQ-9 and the endogenous mouse quinone (Figure S12A). Furthermore, when spiking standards into the kidney lysate, the kidney lysate without any spike in and kidney sample with RQ-9-spiked in had complete overlap, whereas the kidney lysate with the isoRQ-9 spiked in yield a broader and backshifted peak (Figure S12B), confirming that endogenous kidney rhodoquinone is the RQ-9, not isoRQ-9.

Next, we calculated the concentration of RQ-9 in our kidney mitochondria lysates by spiking in purified RQ-9 standard into the kidney sample. Briefly, kidney lysates with an unknown RQ-9 concentration were prepared and split into two identical aliquots. One sample was spiked with our analytical RQ-9 standard to a final concentration of 1 µM, while the other received an equivalent volume of solvent to maintain equal levels of endogenous RQ-9 (Figure 4E). These samples were then analyzed by LC-MS alongside a standard curve of the RQ-9 analytical standard. The ion counts of RQ-9 were determined for all samples as the sum of integrated peak areas from several common adducts including [M+2H]^+^: 781.6372 *m/z*, [M+H]^+^: 780.6294 *m/z*, [M+K]^+^: 818.5853 *m/z*, [M+Na]^+^: 802.6114 *m/z*, and [M+Cu]^+^: 842.5512 *m/z* (Figure 4F). To make sure that the level of RQ-9 in the samples were within the linear detection range of the instrument, the ion counts of the two samples were compared to the standard curve (Figure 4G). Then, the difference in peak areas between the spiked and non-spiked samples were calculated, giving the signal intensity reflective of the 1 µM synthetic RQ-9 standard that we spiked in. Finally, to get the endogenous RQ-9 concentration in the kidney lysate, we applied the response factor reflective of ion counts per 1 µM concentration to the remaining ion counts in the spike-in sample, thereby extrapolating the endogenous RQ-9 concentration. In this case, the concentration of RQ-9 in the kidney lysate was measured to be 1.58 µM. With this method, additional calculations can be performed to obtain the physiological concentration of RQ-9 in a given mitochondrion or even a whole mouse kidney.^4, 27^

## Discussion

In this study, we performed ammonolysis of UQ-9 or UQ-10 to substitute the C-2 or C-3 methoxy group with an amino group, producing a mixture of rhodoquinone (C-2 amino) and isorhodoquinone (C-3 amino). Using flash chromatography, these isomers were separated from one-another, generating analytical-grade standards for use in LC-MS analyses. Future studies may attempt to improve the yield on RQ synthesis by altering the reaction conditions to better favor the ammonia nucleophilic attack at the C-2 position over the C-3 position. Using NMR analysis of RQ-10, this study also confirms with high certainty the positional information of the quinone head group of physiological RQ. Moreover, beyond generating analytical-grade LC-MS standards for RQ, this study also demonstrates the ability of these analytical standards to be converted to the quinol/reduced forms, allowing their chromatographic and fragmentation characterizations by LC-MS/MS. One major application of these analytical RQ standards demonstrated in this study is absolute quantification of the endogenous RQ present in tissue specimen. Beyond absolute quantification, the pure standard is also suitable for reconstitution assays interrogating the biochemical parameters of RQ with mammalian ETCcomplexes, as well as its potential use in structural analyses to map docking sites of RQ on ETC complexes. These purified compounds provide the field necessary reagents to advance future fundamental and translational research on RQ in the mammalian ETC.

## Data Availability

All mass spectrometry raw data files have been deposited on the Metabolomics Workbench repository. All other data are presented in the figures. Additional information is available from the corresponding authors upon request.

## Acknowledgements

We thank Celia Schiffer for her support and guidance on this project. This work was funded by R35GM151996 (C.S. and A.A.), R01 DK144534 (J.B.S.), the Searle Scholars Program (J.B.S.), the Smith Family Awards Program for Excellence in Biomedical Research (J.B.S.), and the Barbara D. Cammett Breakthrough T1D Center of Excellence in New England of the Breakthrough T1D Foundation under award number 4-COE-2025-1751-A-N (J.B.S. & A.A.).

## Author Contributions

T.D. acquired and analyzed LC-MS data, generated and analyzed reduced RQ standards, and performed quantitative analyses of endogenous RQ-9 in mouse tissues. A.A. performed synthesis and purification of standards and acquired and analyzed NMR data. T.D. and J.B.S. wrote the manuscript with edits from A.A. J.B.S. supervised and acquired funding for the study.

## Figure Captions

**Figure S1.** Proton (^1^H) NMR (500 MHz, CDCl_3_) spectra of isorhodoquinone-10.

**Figure S2.** Carbon 13 (^13^C) NMR (126 MHz, CDCl_3_) spectra of isorhodoquinone-10.

**Figure S3.** Distortionless Enhancement by Polarization Transfer (DEPT) NMR (126 MHz, CDCl_3_) spectra of isorhodoquinone-10.

**Figure S4.** Heteronuclear Single Quantum Coherence (HSQC) NMR analysis of isorhodoquinone-10.

**Figure S5.** Heteronuclear Multiple Bond Correlation (HMBC) NMR analysis of isorhodoquinone-10.

**Figure S6.** Proton (^1^H) NMR (500 MHz, CDCl_3_) spectra of rhodoquinone-10.

**Figure S7.** Carbon 13 (^13^C) NMR (126 MHz, CDCl_3_) spectra of rhodoquinone-10.

**Figure S8.** Distortionless Enhancement by Polarization Transfer (DEPT) NMR (126 MHz, CDCl_3_) spectra of rhodoquinone-10.

**Figure S9.** Heteronuclear Single Quantum Coherence (HSQC) NMR analysis of rhodoquinone-10.

**Figure S10.** Heteronuclear Multiple Bond Correlation (HMBC) NMR analysis of rhodoquinone-10.

**Figure S11.** Stability of rhodoquinone-10 standard under light and methanol exposure.

(A) Relative rhodoquinone-10 levels in standard samples diluted in either 80:20 ethanol:hexane or 100% methanol with 2 mM ammonium formate and incubated for 1 hour under dark, ambient light, or ultraviolet (UV) light conditions. Integrated adducts includes [M+H]^+^: 848.6921 *m/z*, [M+Na]^+^: 870.6740 *m/z*, and [M+NH_4_]^+^: 865.7186 *m/z*.

(B) Relative rhodoquinone-10 levels in standards diluted in either 80:20 ethanol:hexane or 100% methanol with 2 mM ammonium formate and incubated for 1 hour under dark, ambient light, or ultraviolet (UV) light conditions. Integrated adducts includes [M+2H]^+^: 851.7155 *m/z*, [M+H]^+^: 850.7077 *m/z*, and [M+Na]^+^: 872.6897 *m/z*.

(C) Relative rhodoquinol-10 to rhodoquinone-10 ratios of standards diluted in either 80:20 ethanol:hexane or 100% methanol with 2 mM ammonium formate and incubated for 1 hour under dark, ambient light, or ultraviolet (UV) light conditions.

(D) Relative ubiquinone-10 generated from resuspending standards in either 80:20 ethanol:hexane or 100% methanol with 2 mM ammonium formate and incubated for 1 hour under dark, ambient light, or ultraviolet (UV) light conditions. Integrated adducts includes [M]^+^: 862.6839 *m/z*, [M+2H]^+^: 864.6996 *m/z*, [M+H]^+^: 863.6917 *m/z*, [M+Na]^+^: 885.6737 *m/z*, and [M+NH_4_]^+^: 880.7183 *m/z*.

For all panels, N = 3 for each condition, values are relative to a same-concentration standard sample without methanolic ammonium formate 2 mM and light exposure (N = 1). Data represent mean ± SEM. P-values were calculated using one-way ANOVA, ns indicates not significant, * p-value < 0.05.

**Figure S12.** C30 Chromatography Analysis of RQ, isoRQ, and mouse tissue.

(A) Overlay of proton adduct chromatograms for endogenous rhodoquinone-9 in mouse kidney, rhodoquinone-9, and isorhodoquinone-9 standards.

(B) Overlay of proton adduct chromatograms for rhodoquinone-9 in mouse kidney sample without standard spiked-in versus with rhodoquinone-9 and isorhodoquinone-9 standards spiked-in.

## Notes

### Competing Interest Statement

The authors have declared no competing interest.

### Summary of Updates

Additional data added looking at stability of RQ standards and added detail into materials and methods section.

## References

(1) Spinelli, J. B.; Haigis, M. C. The multifaceted contributions of mitochondria to cellular metabolism. Nat Cell Biol 2018, 20 (7), 745–754. DOI: 10.1038/s41556-018-0124-1 From NLM.

(2) Martínez-Reyes, I.; Chandel, N. S. Mitochondrial TCA cycle metabolites control physiology and disease. Nat Commun 2020, 11 (1), 102. DOI: 10.1038/s41467-019-13668-3 From NLM.

(3) Van Hellemond, J. J.; Klockiewicz, M.; Gaasenbeek, C. P.; Roos, M. H.; Tielens, A. G. Rhodoquinone and complex II of the electron transport chain in anaerobically functioning eukaryotes. J Biol Chem 1995, 270 (52), 31065–31070. DOI: 10.1074/jbc.270.52.31065 From NLM.

(4) Valeros, J., Jerome, M., Tseyang, T., Vo, P., Do, T., Fajardo Palomino, DC., Grotehans, N., Kunala, M., Jerrett, AE., Hathiramani, NR., Mireku, M., Magesh, RY., Yenilmez, B., Rosen, PC., Mann, JL., Myers, JW., Kunchok, T., Manning, T., Carr, P., Boeker, L., Bin Munim, M., Lewis, CA., Sabatini, DM., Kelly, M., Xie, J., Czech, MP., Gao, G., Shepherd, JN., Walker, AK., Kim, H., Watson, EV., Spinelli, JB. Rhodoquinone Carries Electrons in the mammalian electron transport chain. Cell 2025, 188, 1–16. DOI: 10.1016/j.cell.2024.12.007.

(5) Alberts B, J. A., Lewis J, et al. Garland Science. Electron-Transport Chains and Their Proton Pumps. Molecular Biology of the Cell. 2002, 4th edition.

(6) Spinelli, J. B.; Rosen, P. C.; Sprenger, H. G.; Puszynska, A. M.; Mann, J. L.; Roessler, J. M.; Cangelosi, A. L.; Henne, A.; Condon, K. J.; Zhang, T.;, et al. Fumarate is a terminal electron acceptor in the mammalian electron transport chain. Science 2021, 374 (6572), 1227–1237. DOI: 10.1126/science.abi7495 From NLM.

(7) Guerra, R. M.; Pagliarini, D. J. Coenzyme Q biochemistry and biosynthesis. Trends Biochem Sci 2023, 48 (5), 463–476. DOI: 10.1016/j.tibs.2022.12.006 From NLM.

(8) Kurosu, M.; Begari, E. Vitamin K2 in electron transport system: are enzymes involved in vitamin K2 biosynthesis promising drug targets? Molecules 2010, 15 (3), 1531–1553. DOI: 10.3390/molecules15031531 From NLM.

(9) Lautens, M. J.; Tan, J. H.; Serrat, X.; Del Borrello, S.; Schertzberg, M. R.; Fraser, A. G. Identification of enzymes that have helminth-specific active sites and are required for Rhodoquinone-dependent metabolism as targets for new anthelmintics. PLoS Negl Trop Dis 2021, 15 (11), e0009991. DOI: 10.1371/journal.pntd.0009991 From NLM.

(10) Campbell, A. R. M.; Titus, B. R.; Kuenzi, M. R.; Rodriguez-Perez, F.; Brunsch, A. D. L.; Schroll, M. M.; Owen, M. C.; Cronk, J. D.; Anders, K. R.; Shepherd, J. N. Investigation of candidate genes involved in the rhodoquinone biosynthetic pathway in Rhodospirillum rubrum. PLoS One 2019, 14 (5), e0217281. DOI: 10.1371/journal.pone.0217281 From NLM.

(11) Tan, J. H.; Lautens, M.; Romanelli-Cedrez, L.; Wang, J.; Schertzberg, M. R.; Reinl, S. R.; Davis, R. E.; Shepherd, J. N.; Fraser, A. G.; Salinas, G. Alternative splicing of coq-2 controls the levels of rhodoquinone in animals. Elife 2020, 9. DOI: 10.7554/eLife.56376 From NLM.

(12) Brajcich, B. C.; Iarocci, A. L.; Johnstone, L. A.; Morgan, R. K.; Lonjers, Z. T.; Hotchko, M. J.; Muhs, J. D.; Kieffer, A.; Reynolds, B. J.; Mandel, S. M.;, et al. Evidence that ubiquinone is a required intermediate for rhodoquinone biosynthesis in Rhodospirillum rubrum. J Bacteriol 2010, 192 (2), 436–445. DOI: 10.1128/jb.01040-09 From NLM.

(13) Lester, R. L.; Crane, F. L. The Natural Occurrence of Coenzyme Q and Related Compounds. Journal of Biological Chemistry 1959, 234 (8), 2169–2175. DOI: 10.1016/s0021-9258(18)69886-2.

(14) Pennock, J. F. Occurrence of Vitamins K and Related Quinones Author links open overlay panel. Vitamins and Hormones 1967, 24, 307–329.

(15) Ra., M. Ubiquinones, plastoquinones and vitamins K. Biol Rev Camb Philos Soc. 1971, 1, 47–98.

(16) Castro-Guerrero, N. A.; Jasso-Chávez, R.; Moreno-Sánchez, R. Physiological role of rhodoquinone in Euglena gracilis mitochondria. Biochim Biophys Acta 2005, 1710 (2-3), 113–121. DOI: 10.1016/j.bbabio.2005.10.002 From NLM.

(17) Sato, M., Ozawa, H. Occurrence of Ubiquinone and Rhodoquinone in Parasitic Nematodes, Metastrongylus elongatus and Ascaris lumbricoides var. suis. The Journal of Biochemistry 1969, 65 (6), 861–867.

(18) Allen, P. C. Helminths: Comparison of Their Rhodoquinone. Experimental Parasitology 1973, 34 (2), 211–219.

(19) Takamiya, S., Matsui, T., Taka, H., Murayama, K., Matsuda, M., Aoki, T. Free-Living Nematodes Caenorhabditis elegans Possess in Their Mitochondria an Additional Rhodoquinone, an Essential Component of the Eukaryotic Fumarate Reductase System. ARCHIVES OF BIOCHEMISTRY AND BIOPHYSICS 1999, 371 (2), 284–289.

(20) Van Hellemond, J. J., Luijten, M., Flesch, F.M., Gaasenbeek, C.P.H., Tielens, A.G.M. Rhodoquinone is synthesized de novo by Fasciola hepatica. Molecular and Biochemical Parasitology 1996, 82, 217–226.

(21) James, A. M.; Cochemé, H. M.; Murai, M.; Miyoshi, H.; Murphy, M. P. Complementation of coenzyme Q-deficient yeast by coenzyme Q analogues requires the isoprenoid side chain. Febs j 2010, 277 (9), 2067–2082. DOI: 10.1111/j.1742-4658.2010.07622.x From NLM.

(22) Stefely, J. A.; Pagliarini, D. J. Biochemistry of Mitochondrial Coenzyme Q Biosynthesis. Trends Biochem Sci 2017, 42 (10), 824–843. DOI: 10.1016/j.tibs.2017.06.008 From NLM.

(23) Cluis, C. P.; Burja, A. M.; Martin, V. J. Current prospects for the production of coenzyme Q10 in microbes. Trends Biotechnol 2007, 25 (11), 514–521. DOI: 10.1016/j.tibtech.2007.08.008 From NLM.

(24) Cape, J. L.; Strahan, J. R.; Lenaeus, M. J.; Yuknis, B. A.; Le, T. T.; Shepherd, J. N.; Bowman, M. K.; Kramer, D. M. The respiratory substrate rhodoquinol induces Q-cycle bypass reactions in the yeast cytochrome bc(1) complex: mechanistic and physiological implications. J Biol Chem 2005, 280 (41), 34654–34660. DOI: 10.1074/jbc.M507616200 From NLM.

(25) Moore, H. W., Folkers, K. Coenzyme Q. LXII. Structure and Synthesis of Rhodoquinone, a Natural Aminoquinone of the Coenzyme Q Group. Journal of American Chemical Society 1965, 18 (6), 1409–1410.

(26) Daves, G. D., Jr.; Wilczynski, J. J.; Friis, P.; Folkers, K. Coenzyme Q. CII. Synthesis of rhodoquinone and other multiprenyl 1,4-benzoquinones biosynthetically related to ubiquinone. Journal of the American Chemical Society 1968, 90 (20), 5587–5593. DOI: 10.1021/ja01022a050.

(27) Jerome, M.; Spinelli, J. B. Protocol to extract and measure ubiquinone and rhodoquinone in murine tissues. STAR Protocols 2025, 6 (2). DOI: 10.1016/j.xpro.2025.103880 (accessed 2026/02/23).

(28) Vial, J.; Jardy, A. Experimental Comparison of the Different Approaches To Estimate LOD and LOQ of an HPLC Method. Analytical Chemistry 1999, 71 (14), 2672–2677. DOI: 10.1021/ac981179n.

(29) Burger, N.; Logan, A.; Prime, T. A.; Mottahedin, A.; Caldwell, S. T.; Krieg, T.; Hartley, R. C.; James, A. M.; Murphy, M. P. A sensitive mass spectrometric assay for mitochondrial CoQ pool redox state in vivo. Free Radical Biology and Medicine 2020, 147, 37–47. DOI: 10.1016/j.freeradbiomed.2019.11.028.

(30) Wang, W.; Cao, G.; Zhang, J.; Qiao, H.; Li, H.; Yang, B.; Chen, Y.; Zhu, L.; Sang, Y.; Du, L.;, et al. UV-induced photodegradation of emerging para-phenylenediamine quinones in aqueous environment: Kinetics, products identification and toxicity assessments. J Hazard Mater 2024, 465, 133427. DOI: 10.1016/j.jhazmat.2024.133427.

(31) Ruiz-Jiménez, J.; Priego-Capote, F.; Mata-Granados, J. M.; Quesada, J. M.; Luque de Castro, M. D. Determination of the ubiquinol-10 and ubiquinone-10 (coenzyme Q10) in human serum by liquid chromatography tandem mass spectrometry to evaluate the oxidative stress. J Chromatogr A 2007, 1175 (2), 242–248. DOI: 10.1016/j.chroma.2007.10.055 From NLM.

(32) Xiao, J. F.; Zhou, B.; Ressom, H. W. Metabolite identification and quantitation in LC-MS/MS-based metabolomics. Trends Analyt Chem 2012, 32, 1–14. DOI: 10.1016/j.trac.2011.08.009 From NLM.

(33) Roberts, L. D.; Souza, A. L.; Gerszten, R. E.; Clish, C. B. Targeted metabolomics. Curr Protoc Mol Biol 2012, Chapter 30, Unit 30.32.31–24. DOI: 10.1002/0471142727.mb3002s98 From NLM.

