## Supplemental Figures for "A method to synthesize analytical rhodoquinone standards for quantitative analysis in tissue specimen"

### Figure S1

[1]H NMR ($CDCl_3$, 500 MHz)
Compound 1
isoRhodoquinone-10

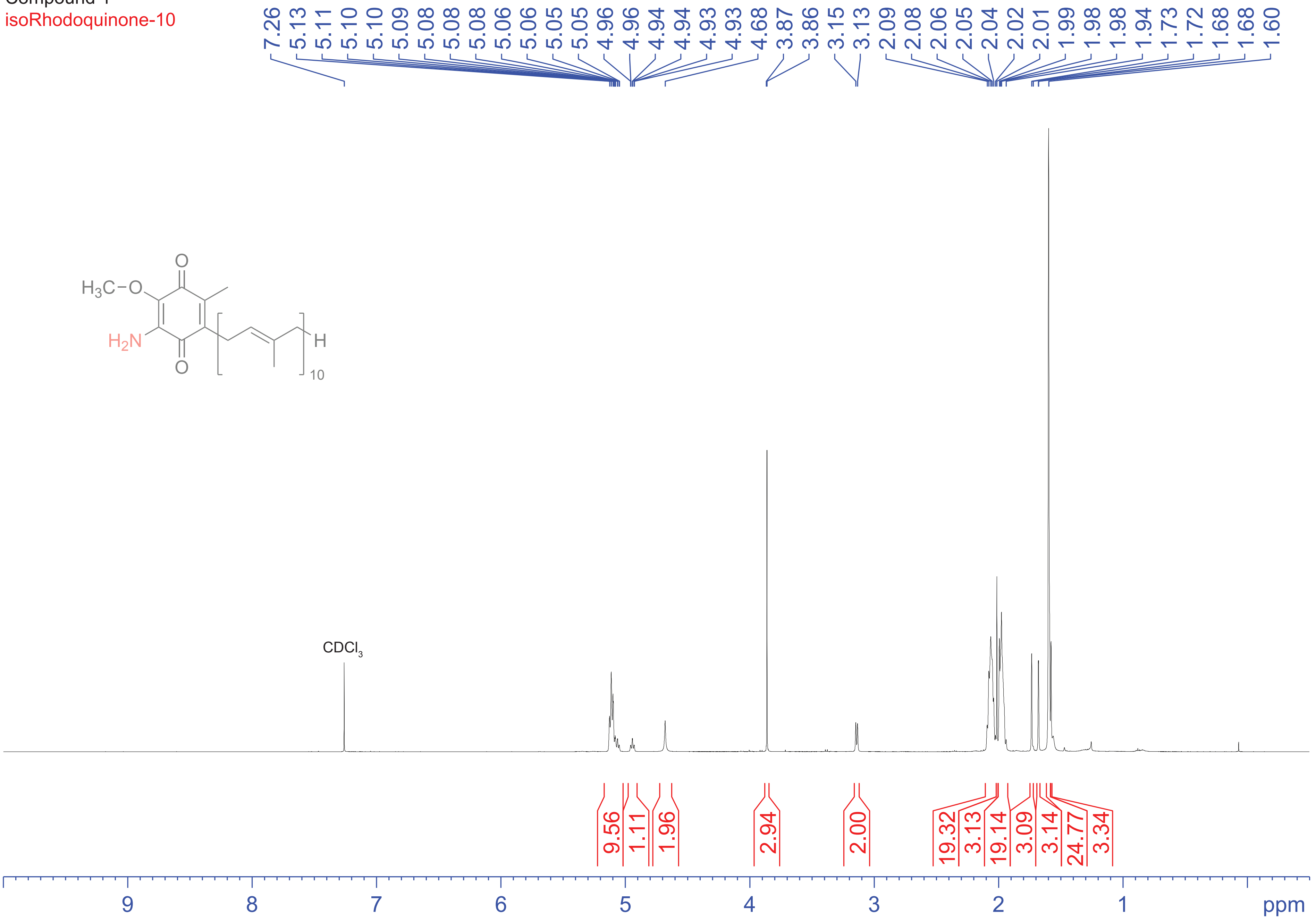

### Figure S2

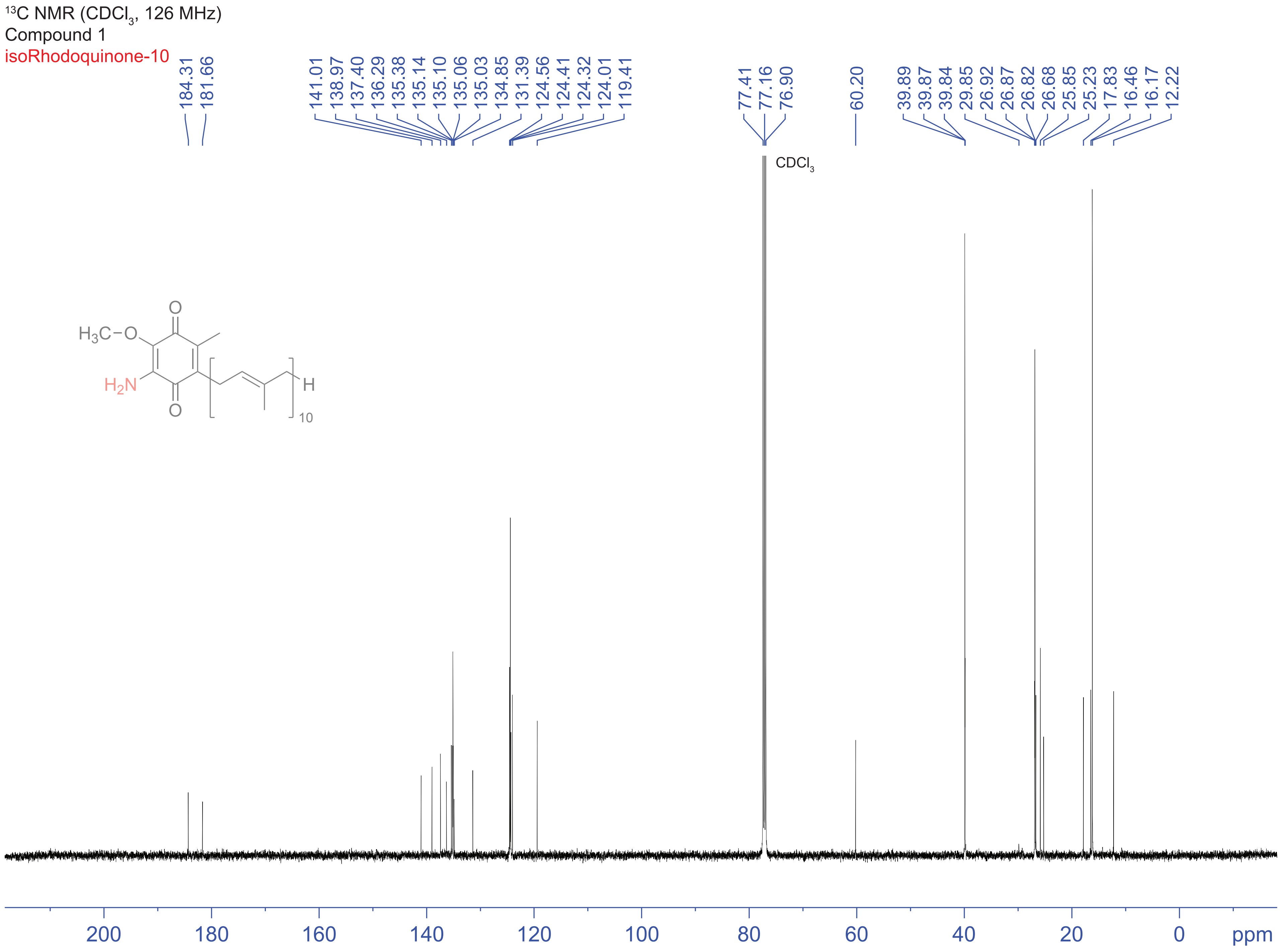

13C NMR (CDCl3, 126 MHz)
Compound 1
isoRhodoquinone-10
184.31
181.66
141.01
138.97
137.40
136.29
135.38
135.14
135.10
135.06
135.03
134.85
131.39
124.56
124.41
124.32
124.01
119.41
77.41
77.16
76.90
60.20
39.89
39.87
39.84
29.85
26.92
26.87
26.82
26.68
25.85
25.23
17.83
16.46
16.17
12.22
CDCl3
O
H3C−O
H2N
O
H
10
200
180
160
140
120
100
80
60
40
20
0
ppm

### Figure S3

DEPT NMR ($CDCl_3$, 126 MHz)
Compound 1
isoRhodoquinone-10

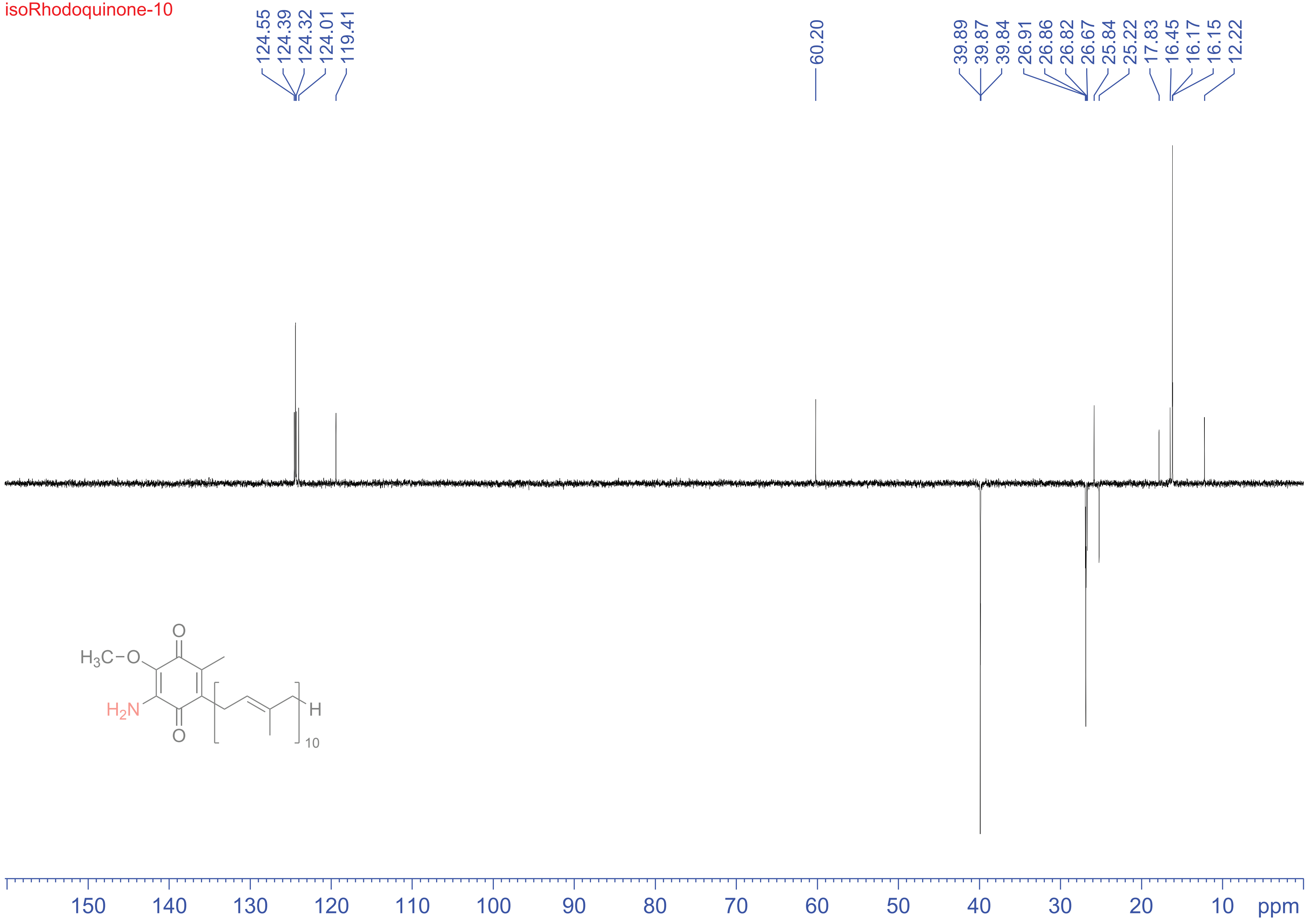

### Figure S4

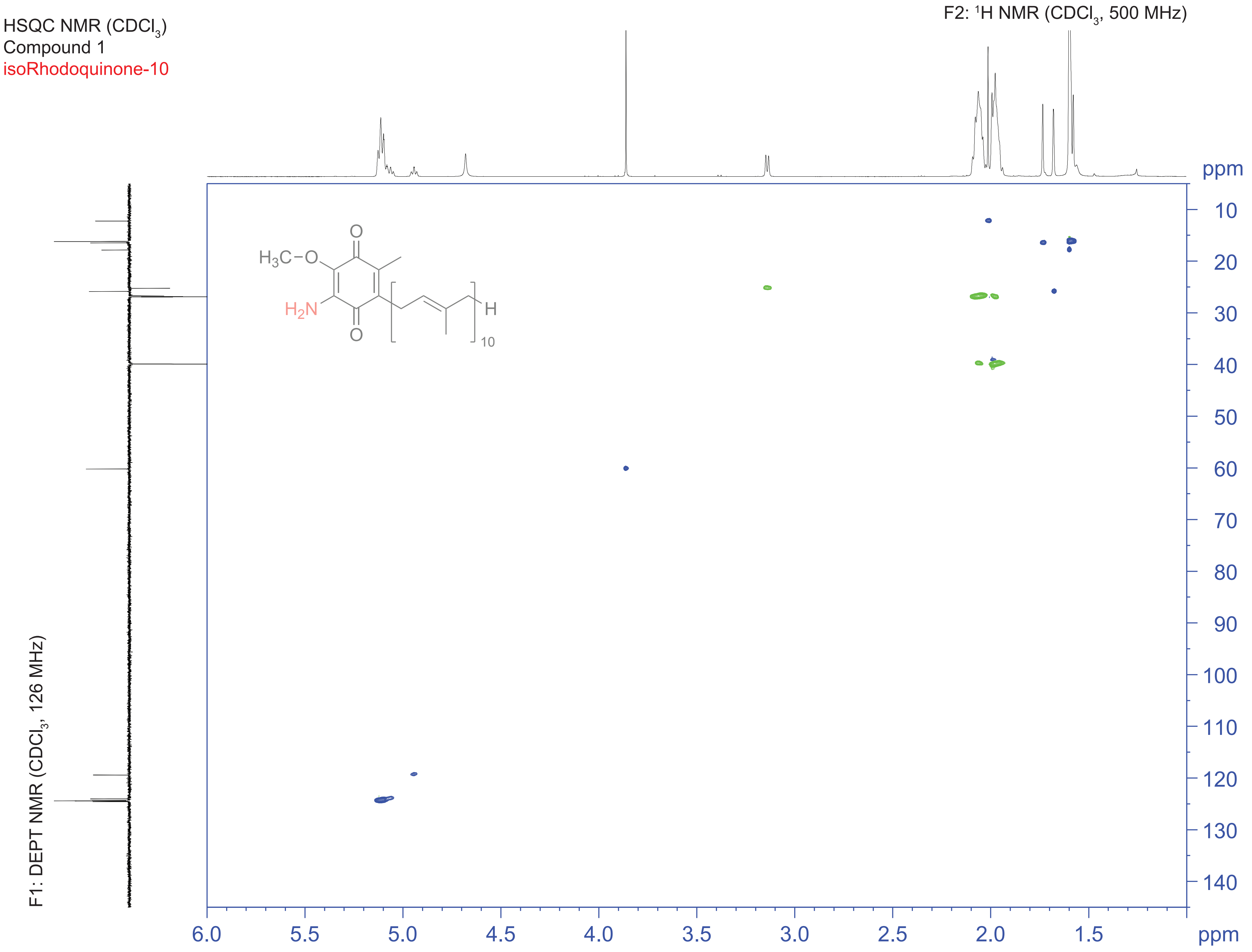

HSQC NMR (CDCl3)
Compound 1
isoRhodoquinone-10
F2: 1H NMR (CDCl3, 500 MHz)
F1: DEPT NMR (CDCl3, 126 MHz)
H3C−O
H2N
O
O
H
10
ppm
10
20
30
40
50
60
70
80
90
100
110
120
130
140
6.0
5.5
5.0
4.5
4.0
3.5
3.0
2.5
2.0
1.5
ppm

### Figure S5

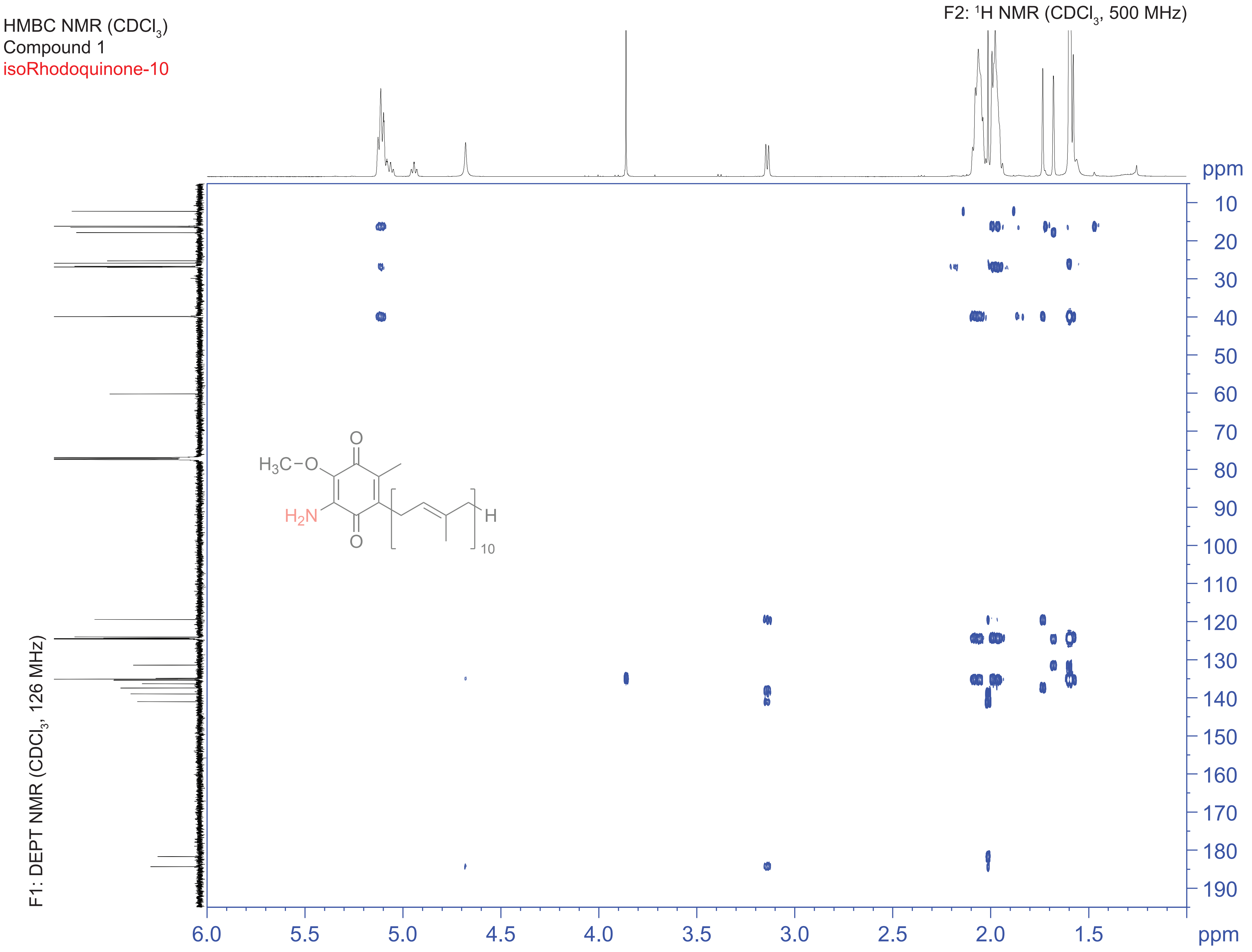
HMBC NMR ($CDCl_3$)
Compound 1
isoRhodoquinone-10
F2: $^1$H NMR ($CDCl_3$, 500 MHz)
F1: DEPT NMR ($CDCl_3$, 126 MHz)
ppm
10
20
30
40
50
60
70
80
90
100
110
120
130
140
150
160
170
180
190
6.0
5.5
5.0
4.5
4.0
3.5
3.0
2.5
2.0
1.5
ppm
$H_3C-O$
$H_2N$
O
O
H
10

### Figure S6

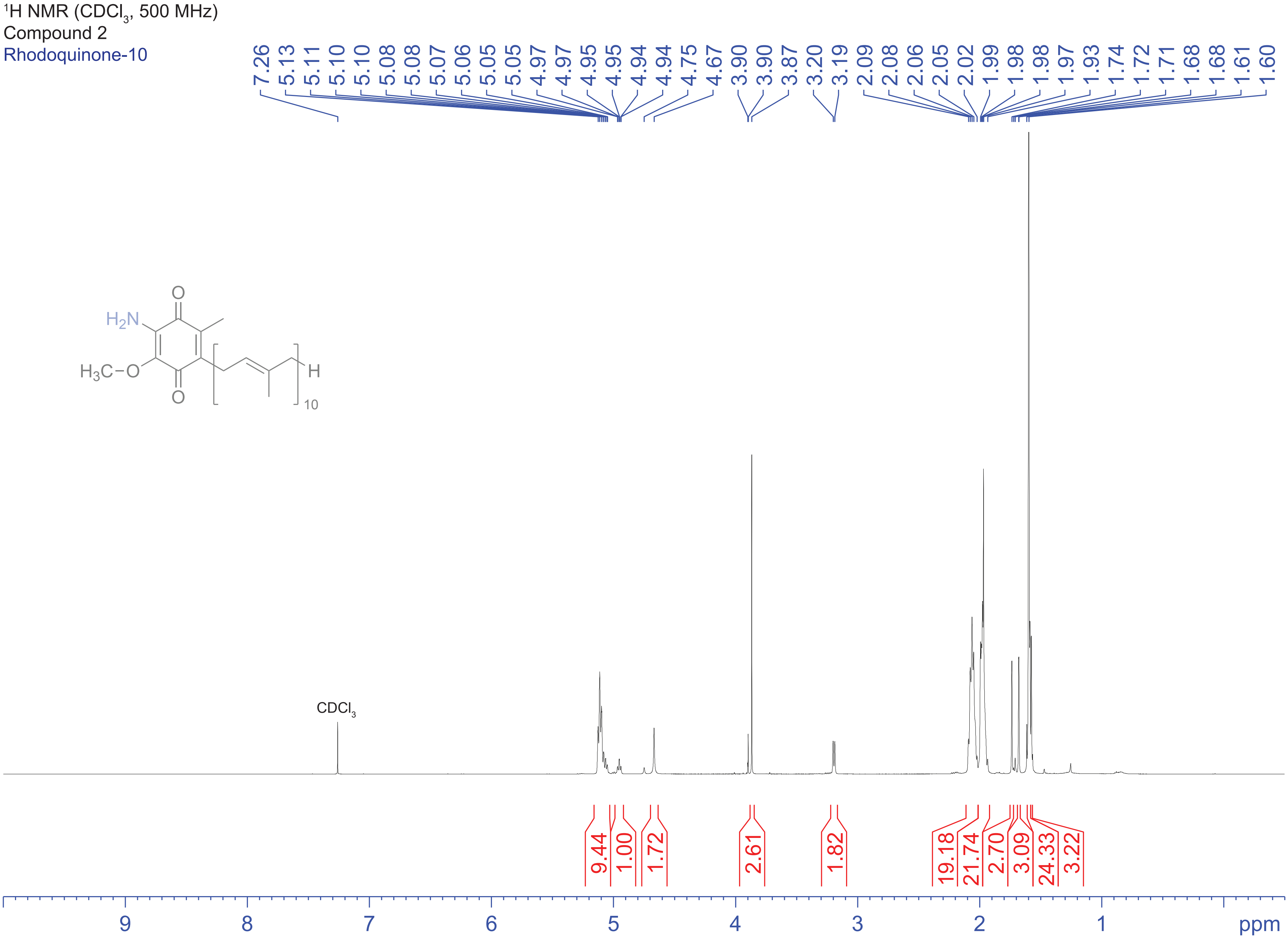

1H NMR (CDCl3, 500 MHz)
Compound 2
Rhodoquinone-10
7.26 5.13 5.11 5.10 5.10 5.08 5.08 5.07 5.06 5.05 5.05 4.97 4.97 4.95 4.95 4.94 4.94 4.75 4.67 3.90 3.90 3.87 3.20 3.19 2.09 2.08 2.06 2.05 2.02 1.99 1.98 1.98 1.97 1.93 1.74 1.72 1.71 1.68 1.68 1.61 1.60
H2N
H3C−O
O
O
H
10
CDCl3
9.44 1.00 1.72 2.61 1.82 19.18 21.74 2.70 3.09 24.33 3.22
9 8 7 6 5 4 3 2 1 ppm

### Figure S7

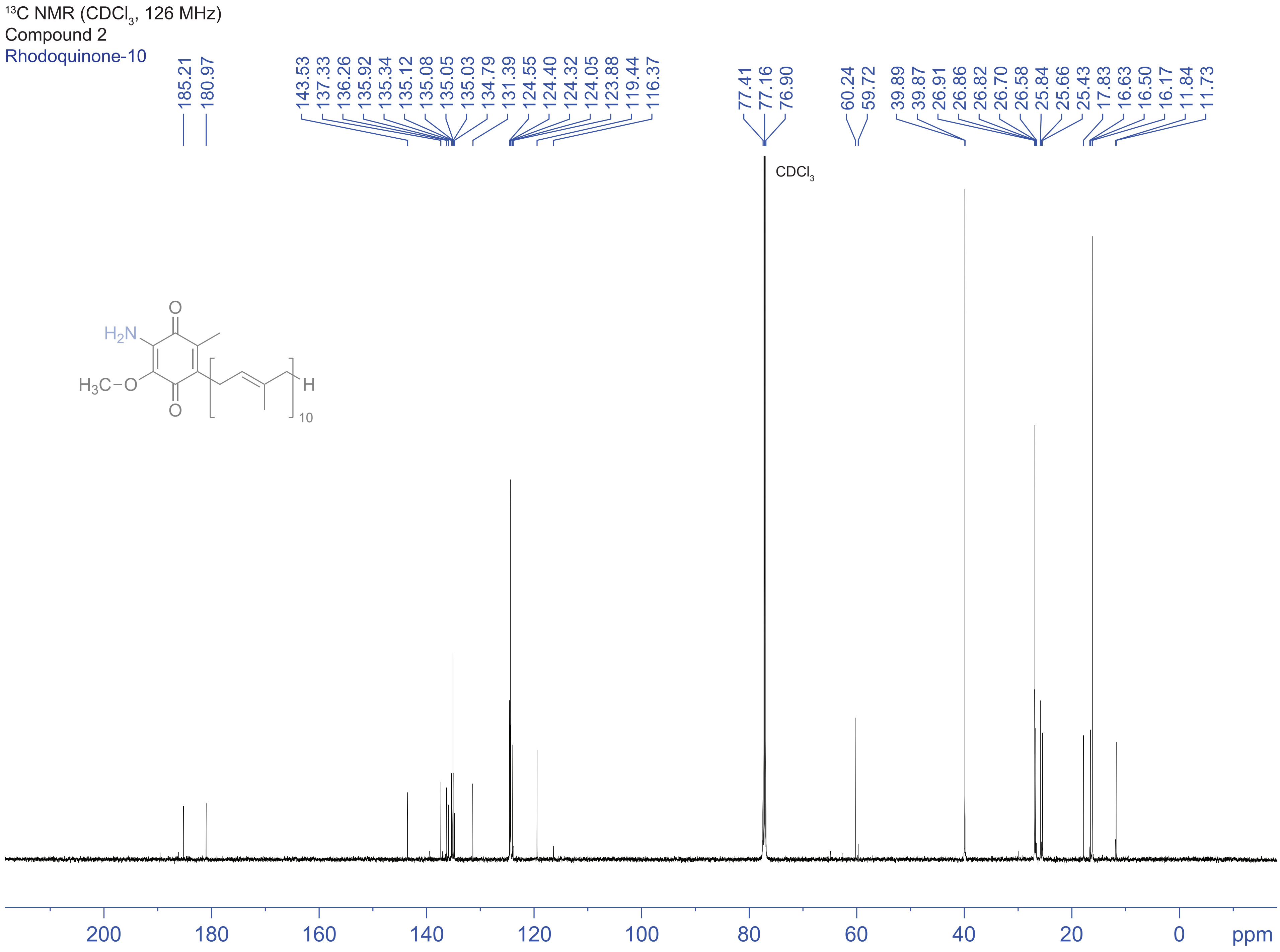

13C NMR (CDCl3, 126 MHz)
Compound 2
Rhodoquinone-10
185.21
180.97
143.53
137.33
136.26
135.92
135.34
135.12
135.08
135.05
135.03
134.79
131.39
124.55
124.40
124.32
124.05
123.88
119.44
116.37
77.41
77.16
76.90
60.24
59.72
39.89
39.87
26.91
26.86
26.82
26.70
26.58
25.84
25.66
25.43
17.83
16.63
16.50
16.17
11.84
11.73
CDCl3
O
H2N
H3C–O
O
H
10
200
180
160
140
120
100
80
60
40
20
0
ppm

### Figure S8

DEPT NMR ($CDCl_3$, 126 MHz)
Compound 2
Rhodoquinone-10

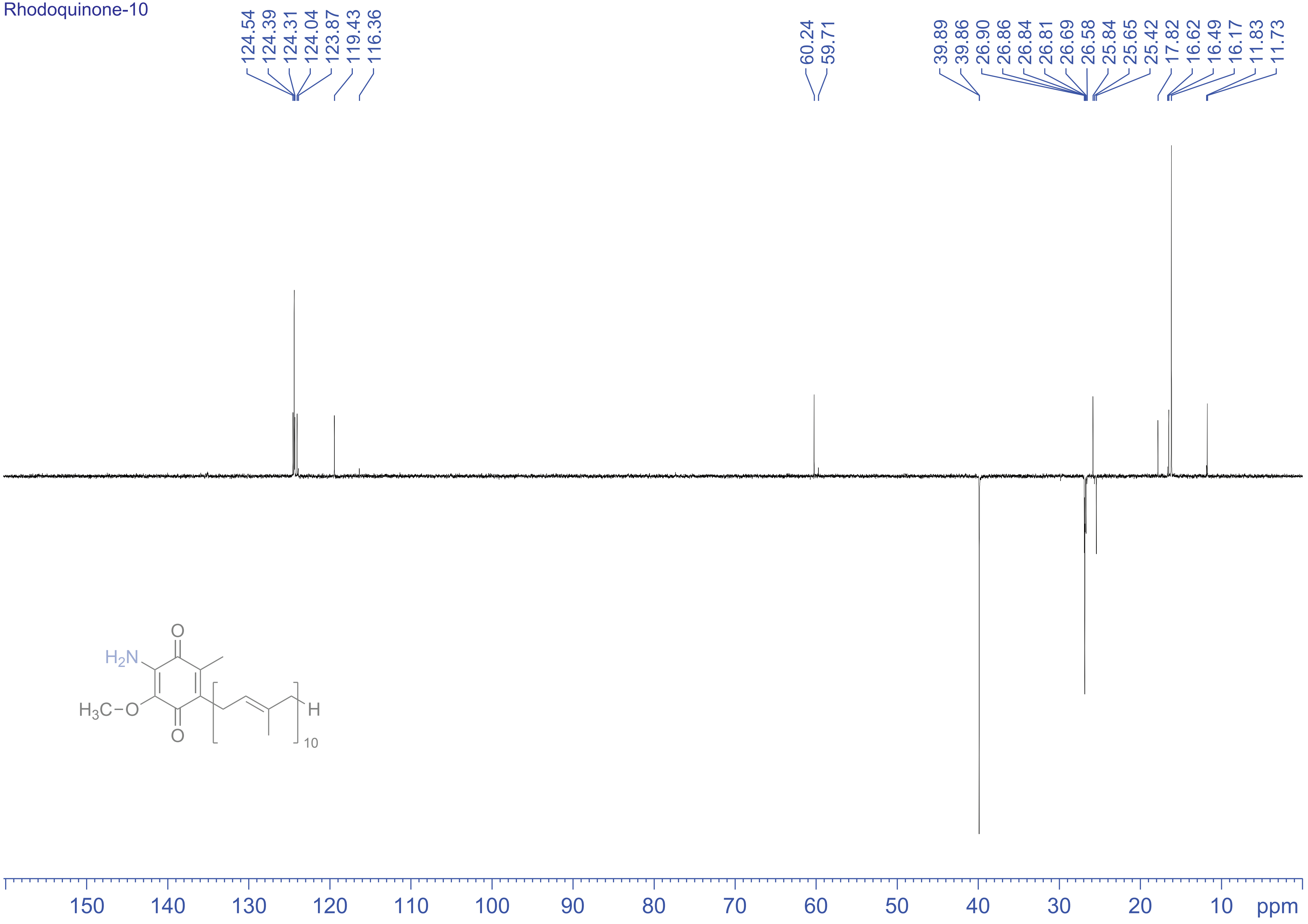

### Figure S9

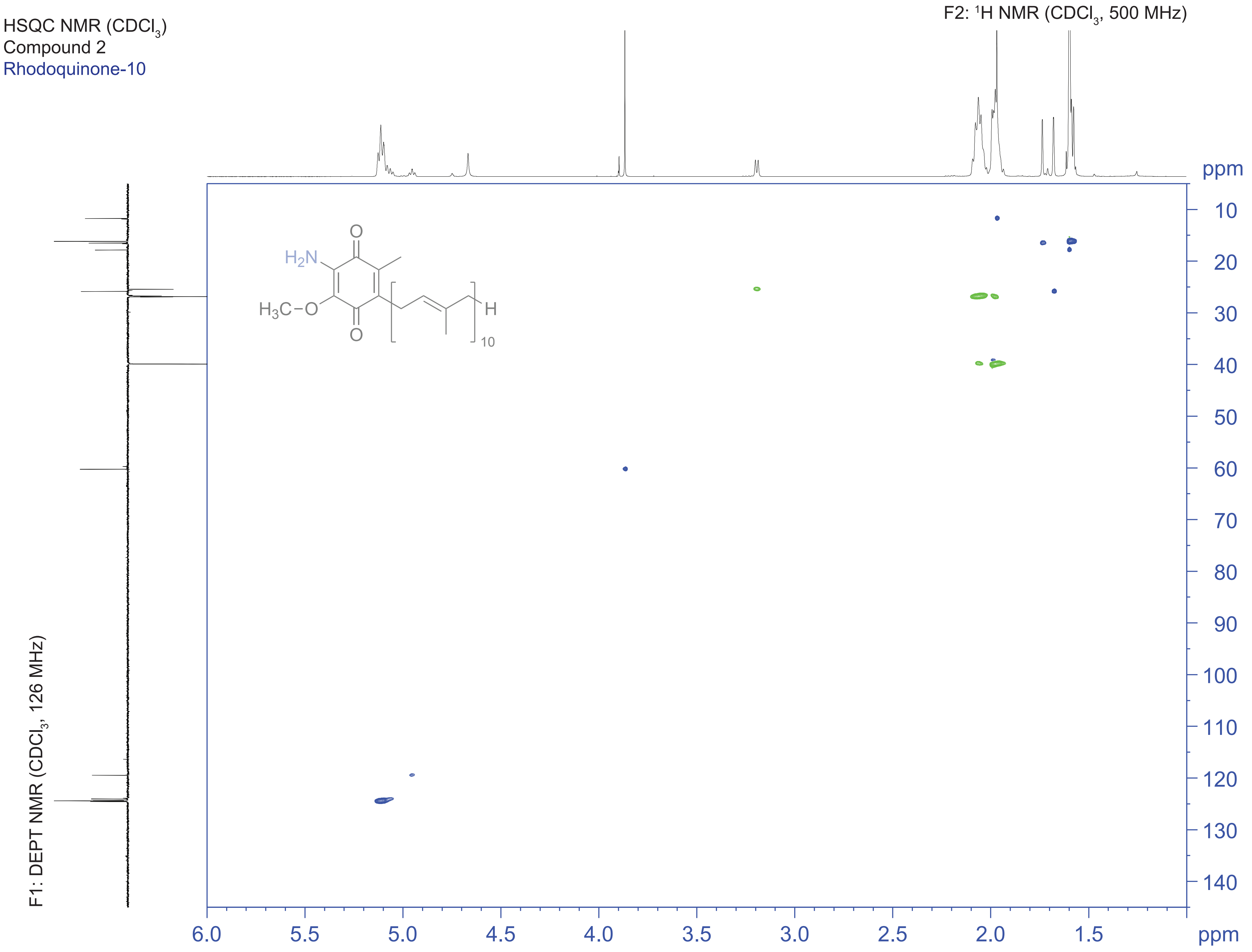

HSQC NMR ($CDCl_3$)
Compound 2
Rhodoquinone-10
F2: $^1H$ NMR ($CDCl_3$, 500 MHz)
F1: DEPT NMR ($CDCl_3$, 126 MHz)
ppm
H2N
H3C–O
O
H
10
6.0
5.5
5.0
4.5
4.0
3.5
3.0
2.5
2.0
1.5
10
20
30
40
50
60
70
80
90
100
110
120
130
140

### Figure S10

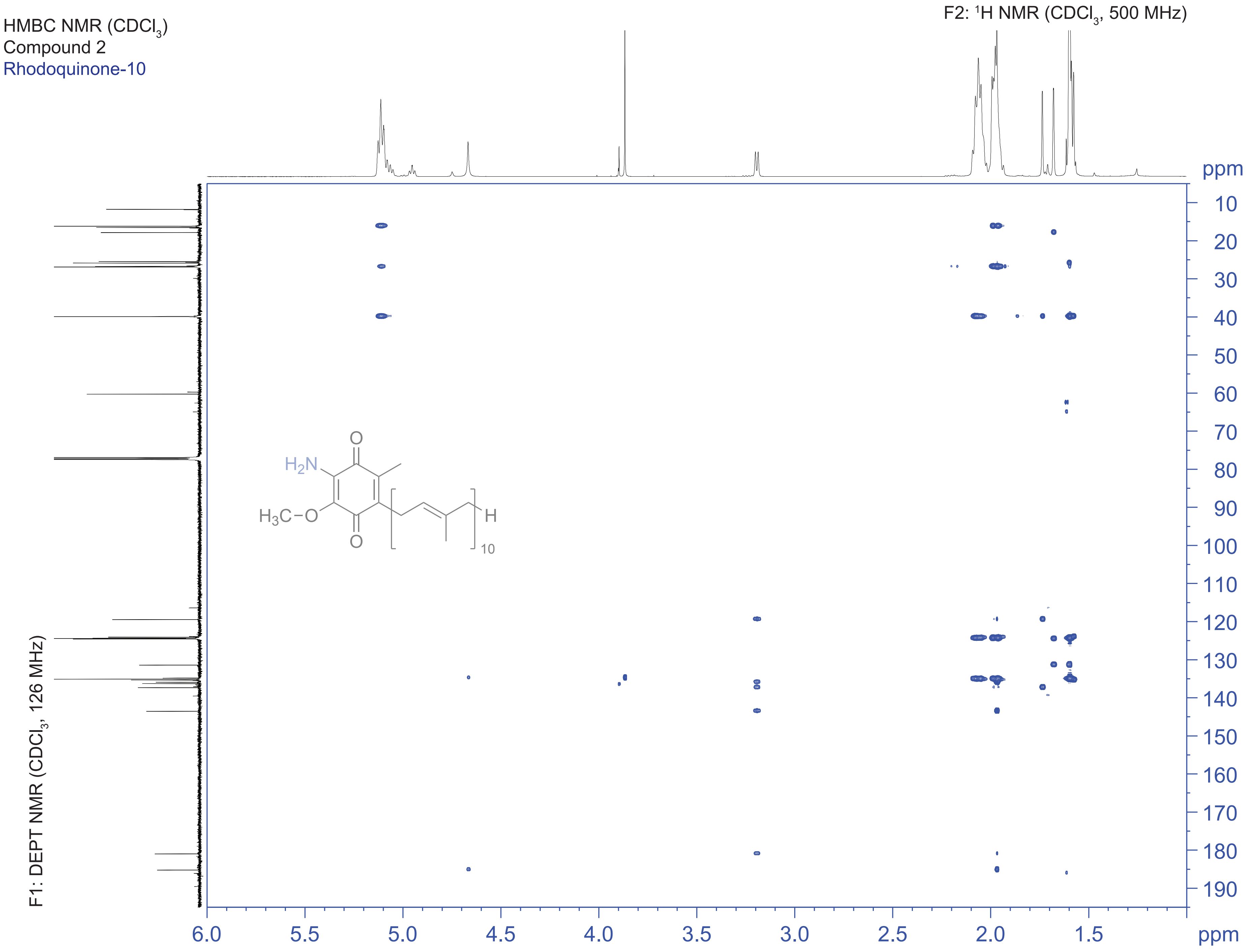

HMBC NMR (CDCl3)
Compound 2
Rhodoquinone-10
F2: 1H NMR (CDCl3, 500 MHz)
F1: DEPT NMR (CDCl3, 126 MHz)
ppm
10
20
30
40
50
60
70
80
90
100
110
120
130
140
150
160
170
180
190
6.0
5.5
5.0
4.5
4.0
3.5
3.0
2.5
2.0
1.5
ppm
O
H2N
H3C–O
O
H
10

### Figure S11

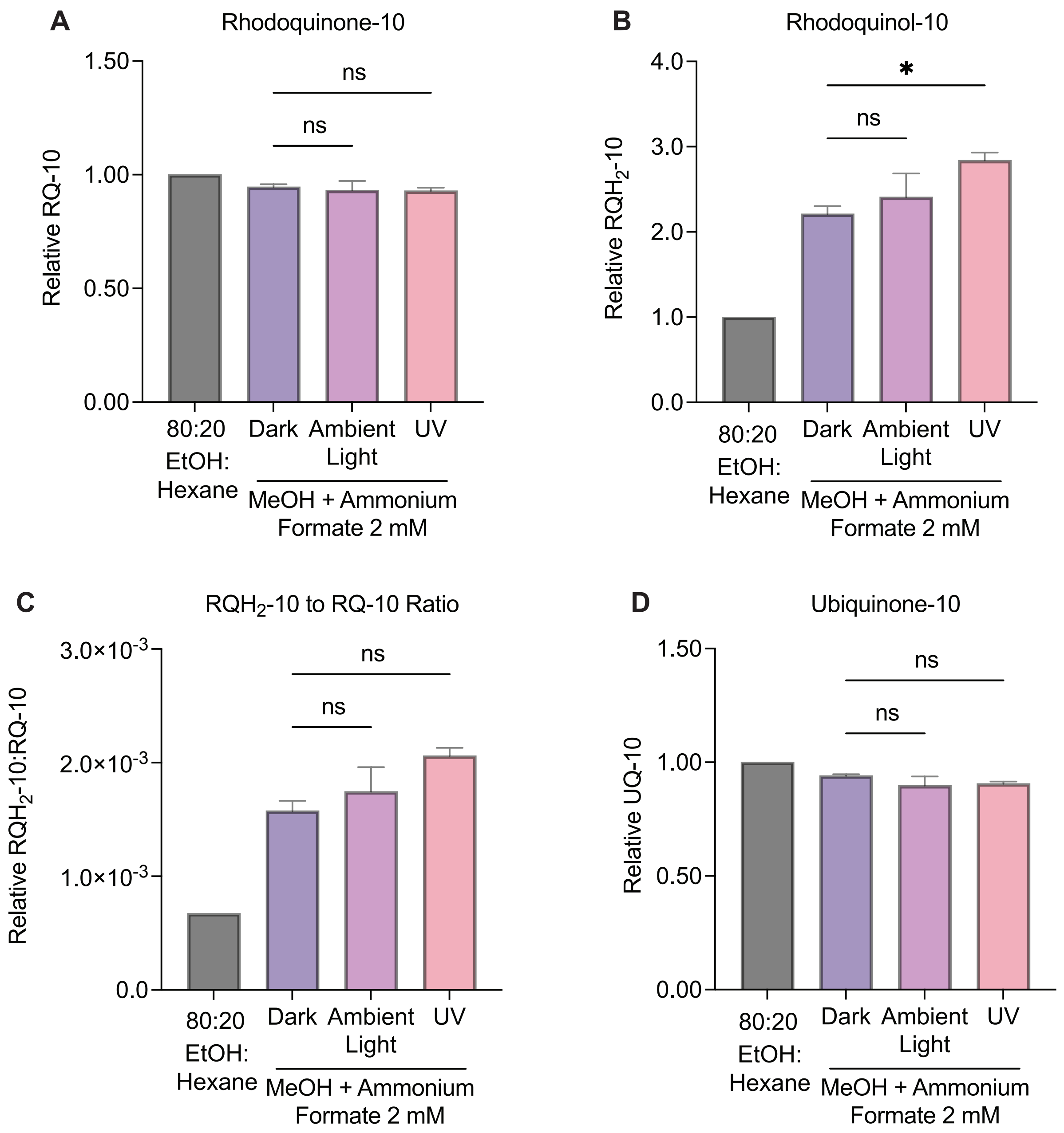
A
Rhodoquinone-10
Relative RQ-10
1.50
1.00
0.50
0.00
ns
ns
80:20 EtOH: Hexane
Dark
Ambient Light
UV
MeOH + Ammonium Formate 2 mM
B
Rhodoquinol-10
Relative $RQH_2$-10
4.0
3.0
2.0
1.0
0.0
*
ns
80:20 EtOH: Hexane
Dark
Ambient Light
UV
MeOH + Ammonium Formate 2 mM
C
$RQH_2$-10 to RQ-10 Ratio
Relative $RQH_2$-10:RQ-10
$3.0\times10^{-3}$
$2.0\times10^{-3}$
$1.0\times10^{-3}$
0.0
ns
ns
80:20 EtOH: Hexane
Dark
Ambient Light
UV
MeOH + Ammonium Formate 2 mM
D
Ubiquinone-10
Relative UQ-10
1.50
1.00
0.50
0.00
ns
ns
80:20 EtOH: Hexane
Dark
Ambient Light
UV
MeOH + Ammonium Formate 2 mM

### Figure S12

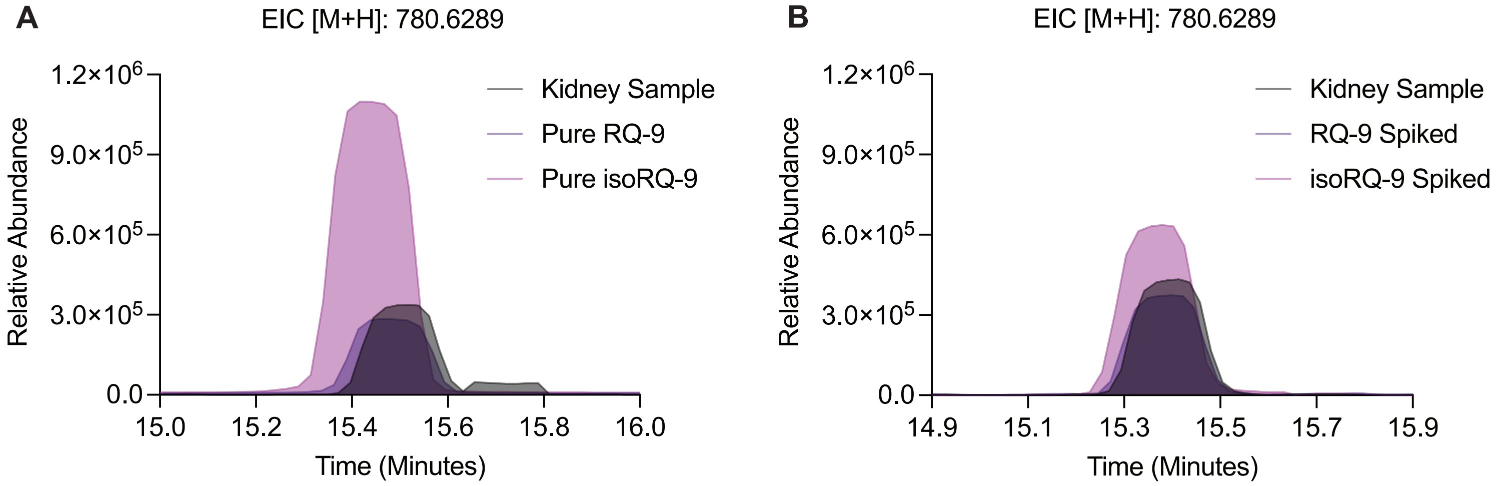
A
EIC [M+H]: 780.6289
Relative Abundance
$1.2\times10^6$
$9.0\times10^5$
$6.0\times10^5$
$3.0\times10^5$
0.0
15.0
15.2
15.4
15.6
15.8
16.0
Time (Minutes)
Kidney Sample
Pure RQ-9
Pure isoRQ-9
B
EIC [M+H]: 780.6289
Relative Abundance
$1.2\times10^6$
$9.0\times10^5$
$6.0\times10^5$
$3.0\times10^5$
0.0
14.9
15.1
15.3
15.5
15.7
15.9
Time (Minutes)
Kidney Sample
RQ-9 Spiked
isoRQ-9 Spiked
